# Targeting cyclic-AMP/phosphodiesterase signaling to quiet nasal tuft cell calcium responses

**DOI:** 10.64898/2026.10.21.760783

**Authors:** Kyle Polen, Li Eon Kuek, Zhongming Ma, Nithin D. Adappa, James N. Palmer, Ryan M. Carey, Robert J. Lee

## Abstract

**Background:** Airway tuft cells, also called solitary chemosensory cells, are enriched in nasal polyps and are likely the main epithelial source of interleukin-25 (IL-25) in chronic rhinosinusitis with nasal polyps (CRSwNP). Tuft cells are activated through G protein-coupled receptors (GPCRs) that signal through Ca^2+^. These receptors may also lower cyclic AMP (cAMP), but whether the cAMP pathway can be used to suppress tuft cell responses is unknown.

**Methods:** Ca^2+^ and cAMP were measured by live cell imaging in isolated Trpm5-GFP mouse nasal tuft cells, and Ca^2+^ responses were measured in primary human nasal epithelial air-liquid interface (ALI) cultures from patients with CRSwNP. Defensin and IL-25 release were measured by ELISA and ciliary beat frequency (CBF) by high-speed imaging. Phosphodiesterase (PDE) expression was assessed using publicly available single cell RNA sequencing datasets.

**Results:** Agonists of bitter taste receptors (T2Rs), the succinate receptor SUCNR1, and the A1 adenosine receptor increased Ca^2+^ in GFP-positive mouse nasal tuft cells. T2R and A1 receptor stimulation also lowered cAMP, and these decreases were blocked by pertussis toxin. Glucose, which activates taste family 1 receptors, increased cAMP. Tuft cells were enriched for *Pde2a*. PDE2A-selective inhibitors (PF-05180999 and BAY 60-7550) increased cAMP and reduced agonist-evoked Ca^2+^ responses in mouse tuft cells. In human nasal ALIs, both compounds reduced denatonium-evoked Ca^2+^ responses, defensin release, and IL-25 release without increasing CBF.

**Conclusions:** GPCRs that activate nasal tuft cells also lower cAMP, and by raising cAMP through PDE2A inhibition, tuft cell Ca^2+^ signaling and IL-25 release were reduced in vitro.

## 1 Introduction

Chronic rhinosinusitis (CRS) and other obstructive airway diseases need better strategies for reducing airway inflammation.^1^ Most current work has focused on monoclonal antibodies that block cytokines or their receptors,^2, 3^ but an alternative approach would be to suppress the epithelial cells upstream in these pathways that release those cytokines. Epithelial-derived IL-25, IL-33, and thymic stromal lymphopoietin (TSLP) initiate and sustain the T helper 2 (Th2) inflammation seen in asthma^4^ and CRS with nasal polyps (CRSwNP).^5^ IL-33 and TSLP come from ciliated, goblet, and basal cells,^2, 6, 7^ but the main epithelial source of IL-25 is likely the airway tuft cell, also called the solitary chemosensory cell or brush cell.^8^

Tuft cells are key mediators of Th2 inflammation in the gut^9–13^ and likely also in the airway. In mice, nasal and tracheal tuft cells release acetylcholine (ACh), which regulates breathing,^14, 15^ inflammation,^16^ ciliary beating, and mucociliary clearance.^17, 18^ Cysteinyl leukotrienes, possibly downstream of purinergic receptors, may regulate expansion of tuft cells during Th2 inflammation,^19, 20^ the phenotype of most CRSwNP and asthma. Tuft cell IL-25 and leukotrienes may act together to perpetuate that inflammation.^21^ Tuft cells may also be important in viral infection,^22^ appearing in the distal lung after severe infection.^23, 24^ In humans, electron micrograph studies suggest that airway tuft cells increase in diseases such as primary ciliary dyskinesia.^25–29^ Tuft cells are enriched in human nasal polyps^30, 31^ and may be the primary epithelial source of IL-25 in CRSwNP.^32^ Because lowering IL-25 may benefit patients with CRSwNP,^33^ quieting tuft cells may be a reasonable target to reduce CRS inflammation.

Sinonasal tuft cells express “taste” signaling components similar to type II cells of the taste bud,^15, 34–36^ including G protein-coupled receptors (GPCRs) for bitter taste (taste family 2 receptors, or T2Rs), the taste-related Gα subunit gustducin (Gα-gustducin), and the ion channel TRPM5. Tuft cells likely use signaling similar to type II taste cells (**Figure 1**),^37^ but rather than purinergic transmission to gustatory neurons, the end result in the nose is secretion of IL-25 and/or ACh through mechanisms that are not yet defined. Tuft cell T2Rs likely detect bacterial and possibly fungal metabolites. ^15, 38–40^ Tuft cells may also respond to odorant irritants^41^ or other host signals via GPCRs.

**Figure 1.**
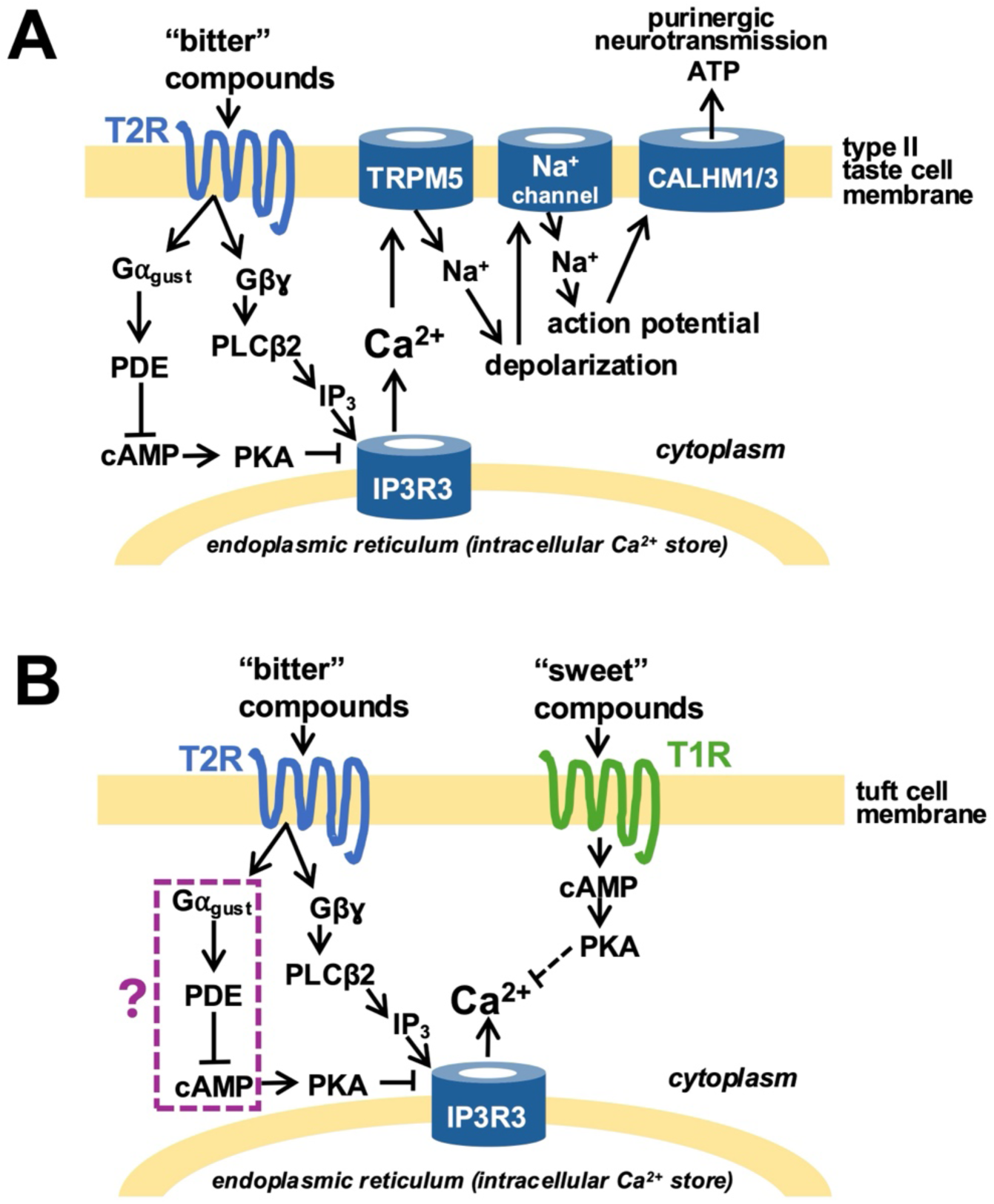
Signaling pathways in type II taste cells and nasal tuft cells. **(A)** Nasal tuft cells have been proposed to use pathways similar to those of type II taste cells,^37^ which use GPCRs to detect bitter, sweet, or umami tastes. Activation of taste GPCRs leads to a Gβγ-mediated increase in Ca^2+^ and a Gα-mediated decrease in cAMP that enhances the Ca^2+^ signal. **(B)** In human nasal tuft cells, we previously showed that T2R activation enhances Ca^2+^ signaling while T1R activation by sugars^42^ or bacterial D-amino acids^43^ represses these responses. We hypothesized that this occurred through cAMP/PKA signaling. Our goal here was to test more directly whether T2Rs and other Ca^2+^-coupled GPCRs in tuft cells reduce cAMP, and whether this affects Ca^2+^ signaling. Abbreviations: CALHM1/3, calcium homeostasis modulator 1/3; cAMP, cyclic adenosine monophosphate; IP_3_, inositol trisphosphate; IP_3_R_3_, type III IP_3_ receptor; PLCβ2, phospholipase C isoform β2; PDE, phosphodiesterase; PKA, protein kinase A; T1R, taste family 1 sweet/umami receptor; T2R, taste family 2 bitter receptor; TRPM5, transient receptor potential melastatin 5 ion channel.

We previously found that human nasal tuft cells respond to T2R agonists by triggering antimicrobial peptide secretion from surrounding epithelial cells at levels sufficient to kill airway pathogens including *Pseudomonas aeruginosa* and *Staphylococcus aureus*.^42^ Those responses were driven by Ca^2+^ signals, likely downstream of T2R activation of Gα-gustducin, which is a marker of nasal tuft cells.^36^ Stimulating tuft cell taste family 1 sweet receptors (T1Rs) with airway surface liquid sugars^42^ or “sweet” bacterial D-amino acids^43^ suppressed the same responses, which we hypothesized occurred through cAMP signaling. T2Rs and T1Rs both couple to gustducin on the tongue,^44–46^ but T1Rs have also been reported to modulate Gαs-cAMP signaling in other cell types, possibly by Gαs coupling.^47–51^

We hypothesize that we might be able to co-opt the cAMP pathway to reduce tuft cell signaling to perhaps reduce inflammation in CRSwNP or other inflammatory airway diseases. Nasal tuft cells express T2Rs^15, 35, 36, 41–43, 52^ and T1Rs,^42, 43, 53^ and possibly formyl peptide receptors,^17^ SUCNR1/GPR91 succinate receptors,^54^ and others.^55^ Rather than targeting any one receptor, targeting the shared cAMP and Ca^2+^ machinery might offer a more robust approach. Gustducin is the Gα subunit for many of these receptors. It is closely related to the two retinal transducin isoforms and pairs with Gβ3 and Gγ13 in taste cells^56^ to generate inositol trisphosphate (IP_3_) through phospholipase C isoform β2 (PLCβ2),^56^ which in turn activates the type III IP_3_ receptor (IP_3_R).^57^ Gustducin itself is thought to lower cAMP by activating phosphodiesterases (PDEs).^58^ That fall in cAMP has been proposed to relieve cAMP/protein kinase A (PKA) repression of either the type III IP_3_R or PLCβ2.^59–61^ In tongue taste cells, tonic gustducin activity has been proposed to hold cAMP low and thereby enhance Ca^2+^ responses to tastants;^59^ however, the full implications of gustducin signaling are not fully elucidated. In particular, the type III IP_3_R has at least three PKA phosphorylation sites,^62^ and PKA has been reported both to stimulate^63^ and to inhibit^61^ this receptor.

Our goal was to test whether the cAMP pathway can be used to reduce nasal tuft cell activation. We used primary mouse and human nasal epithelial cells to further elucidate the role of cAMP and PDEs in tuft cell Ca^2+^ signaling and downstream responses using a combination of live cell imaging and biochemical assays. We also used gene expression insights from publicly available single cell RNAseq databases to better understand how we might be able to target tuft cells in CRSwNP or other inflammatory airway diseases.

## 2 Materials and Methods

### 2.1 Animals

Nasal septa were taken from Trpm5-GFP mice on a C57BL/6 background^64^ that had been euthanized for unrelated experiments under protocols approved by the Institutional Animal Care and Use Committees of the University of Pennsylvania. No animals were sacrificed for the purposes of this study.

### 2.2 Human subjects

Residual surgical tissue was obtained under University of Pennsylvania institutional review board protocol #800614 from adults having sinonasal surgery for CRS. Every donor provided written informed consent. The work followed institutional policy governing the use of residual clinical material, 45 CFR 46.116, and the Declaration of Helsinki. Donors had to be at least 18 years of age. We excluded anyone who had received antibiotics, oral corticosteroids, or a biologic agent such as omalizumab within the month before surgery, anyone with a systemic disease such as granulomatosis with polyangiitis, cystic fibrosis, or a systemic immunodeficiency, and pregnant or cognitively impaired individuals.

### 2.3 Data analysis and statistics

Comparisons between two groups used unpaired two-tailed t-tests. Comparisons among more than two groups used one-way ANOVA with Dunnett’s, Bonferroni, or Tukey’s post test as indicated in each figure legend. All analyses were performed in GraphPad Prism version 10 (San Diego, CA, USA). Figures are marked *p* < 0.05 (*), *p* < 0.01 (**), *p* < 0.001 (***), and *p* < 0.0001 (****); comparisons that were not significant are labeled N.S. or left unmarked. Data are mean ± SEM of at least three independent experiments. Experiments using primary human cells were repeated in cells from at least three separate patients. Single cell gene expression data were analyzed and plotted directly on the Broad Institute Single Cell Portal (https://singlecell.broadinstitute.org/), with the individual datasets referenced and linked in the text and figure legends.

### 2.4 Additional methods

Mouse nasal air-liquid interface (ALI) cultures,^65^ human nasal epithelial cell isolation and culture,^66^ live cell imaging,^67,68^ and ciliary beat frequency analysis^69^ followed previously described methods. Full details, including solution compositions and reagent sources and concentrations, are given in the Supporting Information.

## 3 Results

### 3.1 Trpm5-GFP identifies mouse nasal tuft cells that express canonical taste signaling components

In mice, airway and gut tuft cells can be labeled using transgenic animals expressing GFP under a *Trpm5* promoter (Trpm5-GFP).^64, 70, 71^ TRPM5 is part of the canonical taste signaling pathway (**Figure 1**), and while its full role in tuft cells is still unclear, its importance is supported by observations that *Trpm5*^-/-^ mice are more susceptible to airway infection.^72^ We visualized Trpm5-GFP tuft cells in ALI cultures of mouse nasal septum (**Figure 2A**). Tuft cell numbers appeared relatively stable across 2 to 3 weeks of ALI differentiation (**Figure 2B**) but increased with IL-13 treatment (**Figure 2C-D**), as expected.^31, 32, 73, 74^ Analysis of a publicly available single cell gene expression dataset that includes mouse tracheal tuft cells^55^ supported the identity of these Trpm5-expressing cells, based on enrichment of the tuft cell markers *Pou2f3*,^20, 75^ *Dclk1*,^76^ *Avil*,^21, 76^ *Il25*,^32^ and choline acetyltransferase (*Chat*)^77, 78^ (**Figure 2E-F, Supplemental Figure 1**). Mouse tracheal tuft cells in this dataset also expressed classic taste receptor signaling components Gα-gustducin (*Gnat3*), Gβ3 (*Gnb3*), Gγ13 (*Gng13*), *Plcb2*, and the type III IP_3_R (**Figure 2F, Supplemental Figure 2**).^34, 79^ Gustducin appeared to be restricted to tuft cells, but tuft cells expressed a range of Gα isoforms, including Gαs (**Supplemental Figure 3**).

**Figure 2.**
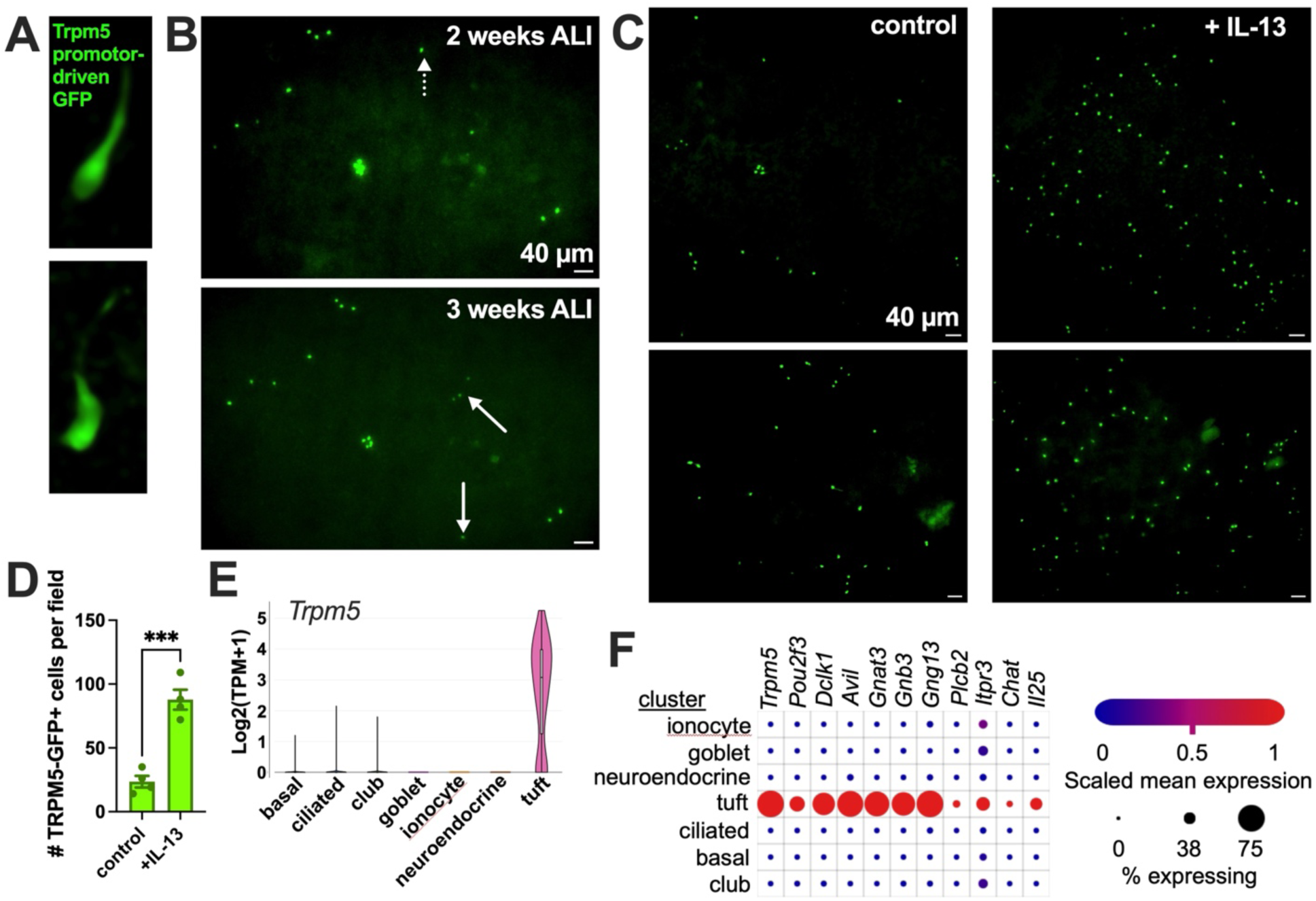
Trpm5-GFP labels tuft cells in mouse nasal air-liquid interface cultures. **(A)** Image of a mouse nasal solitary chemosensory cell grown in air-liquid interface (ALI) culture, as described.^65^ **(B)** Images of nasal ALIs at lower magnification showing the tuft cell population in the same field at 2 and 3 weeks of ALI differentiation. Dotted arrow denotes tuft cell visible at 2 weeks but not 3 weeks. Solid arrows indicate tuft cells visible at 3 weeks that were not visible at 2 weeks. **(C)** Mouse nasal ALIs without (left) or with (right) IL-13 treatment as described in the Supporting Information. **(D)** Quantification of nasal tuft cells per field from experiments as in C. **(E)** Data from Montoro et al.,^55^ showing expression of *Trpm5* in cells identified as tuft cells from mouse trachea. **(F)** Dot plot of data from Montoro et al.,^55^ showing expression of tuft cell markers in mouse tracheal tuft cells. Significance in D by Student’s *t* test; \*\*\**p* < 0.001. Data for E and F were accessed and plotted through the Broad Institute Single Cell Portal (https://singlecell.broadinstitute.org/single_cell/study/SCP163/airway-epithelium).

### 3.2 T2R, SUCNR1, and A1 adenosine receptor agonists activate Ca^2+^ signals in nasal tuft cells

We measured Ca^2+^ in Trpm5-GFP-positive cells acutely isolated from mouse nasal septum. Only GFP-positive cells responded to the T2R agonist denatonium benzoate^42^ (**Figure 3A**), and these responses were eliminated after pre-incubation with PLC inhibitor U73122 but not inactive analogue U73343 (**Figure 3B**). GFP-positive cells also responded to the SUCNR1 agonist succinate (**Figure 3C**). SUCNR1 is another GPCR expressed in nasal tuft cells (**Supplemental Figure 4**).^54, 55^ Because a prior study suggested A1 adenosine receptor (A1AR, encoded by *Adora1*) expression in tuft cells^55^ (**Supplemental Figure 4**), we tested the A1AR agonist N^6^-cyclopentyladenosine (CPA). CPA increased Ca^2+^ in GFP-positive cells (**Figure 3D**), and these responses were blocked by U73122 (**Figure 3E**) and by pertussis toxin (PTX; **Figure 3F**), which ADP-ribosylates gustducin and inhibits signaling of the G protein complex.^80^ The concentration-response relationship for CPA is shown in **Figure 3G**.

**Figure 3.**
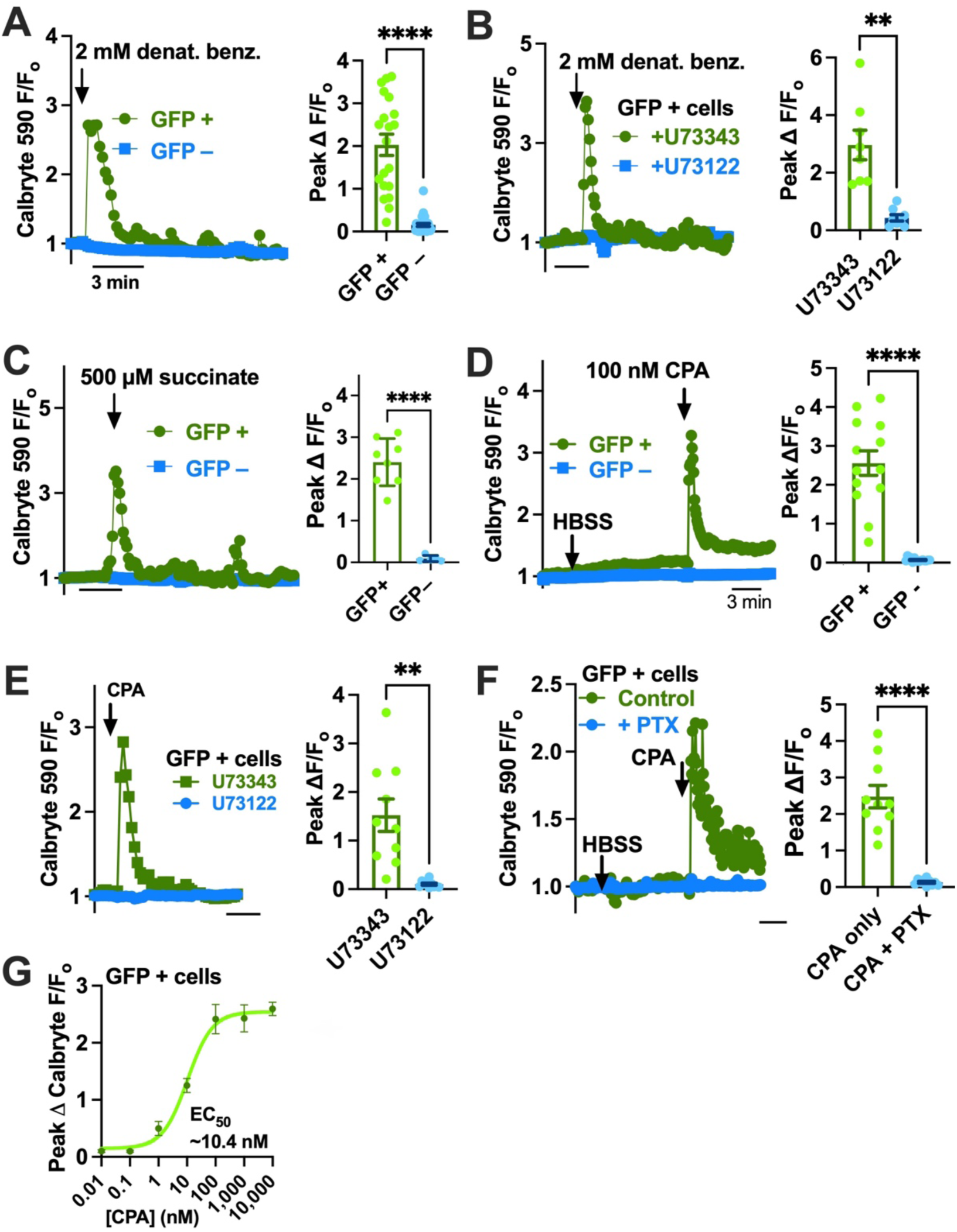
GPCR-stimulated Ca^2+^ signals in Trpm5-GFP-positive mouse nasal tuft cells. Ca^2+^ signals were recorded from freshly isolated mouse nasal tuft cells as described in the Supporting Information. **(A-B)** Representative traces and quantification showing activation of Ca^2+^ responses by the T2R agonist denatonium benzoate in GFP-positive cells (A) and inhibition by the PLC inhibitor U73122 (B). **(C)** The SUCNR1 agonist succinate likewise increased Ca^2+^ only in Trpm5-GFP-positive cells. **(D-F)** The A1 adenosine receptor agonist CPA increased Ca^2+^ in Trpm5-GFP-positive cells (D), and this was inhibited by U73122 (E) or PTX (F). **(G)** Concentration-response relationship for CPA in Trpm5-GFP-positive cells. Significance in A-F by Student’s *t* test; \*\**p* < 0.01, \*\*\*\**p* < 0.0001.

### 3.3 T2R and A1 adenosine receptors lower cAMP, while T1R2/3 activation raises it

To better understand how cAMP changes during receptor stimulation, we isolated nasal septal tuft cells, immediately transduced them with a BacMam expressing a cAMP indicator (red upward cADDis), and plated them on Matrigel for 24 hours. Cells showing both green (GFP) and red (cADDis) fluorescence were then imaged during stimulation with denatonium benzoate or vehicle, with or without PTX (**Figure 4A-C**). The adenylyl cyclase activator forskolin (10 µM) was used as a control to raise cAMP. Denatonium benzoate lowered cAMP (**Figure 4B**), and PTX blocked this effect (**Figure 4C**). The A1AR agonists CPA, adenosine monophosphate (AMP), and adenosine also lowered cAMP (**Figure 4D**). PTX blocked the decreases produced by CPA and adenosine (**Figure 4E**), and the A1AR antagonist 8-cyclopentyltheophylline (CPT) blocked the decrease produced by adenosine (**Figure 4F**). Maximal decreases after 5 min of stimulation are summarized in **Figure 4G**. The maximal increase in cAMP produced by subsequent forskolin application was also reduced by T2R or A1AR stimulation (**Figure 4H**). These data suggest that these receptors engage G protein signaling that reduces steady-state cAMP, either through Gα_i_ inhibition of basal or stimulated adenylyl cyclase activity or through gustducin activation of PDE activity. In contrast, glucose (5 mM), which activates T1R2/3 in nasal tuft cells,^42, 43^ increased cAMP (**Figure 4I**).

**Figure 4.**
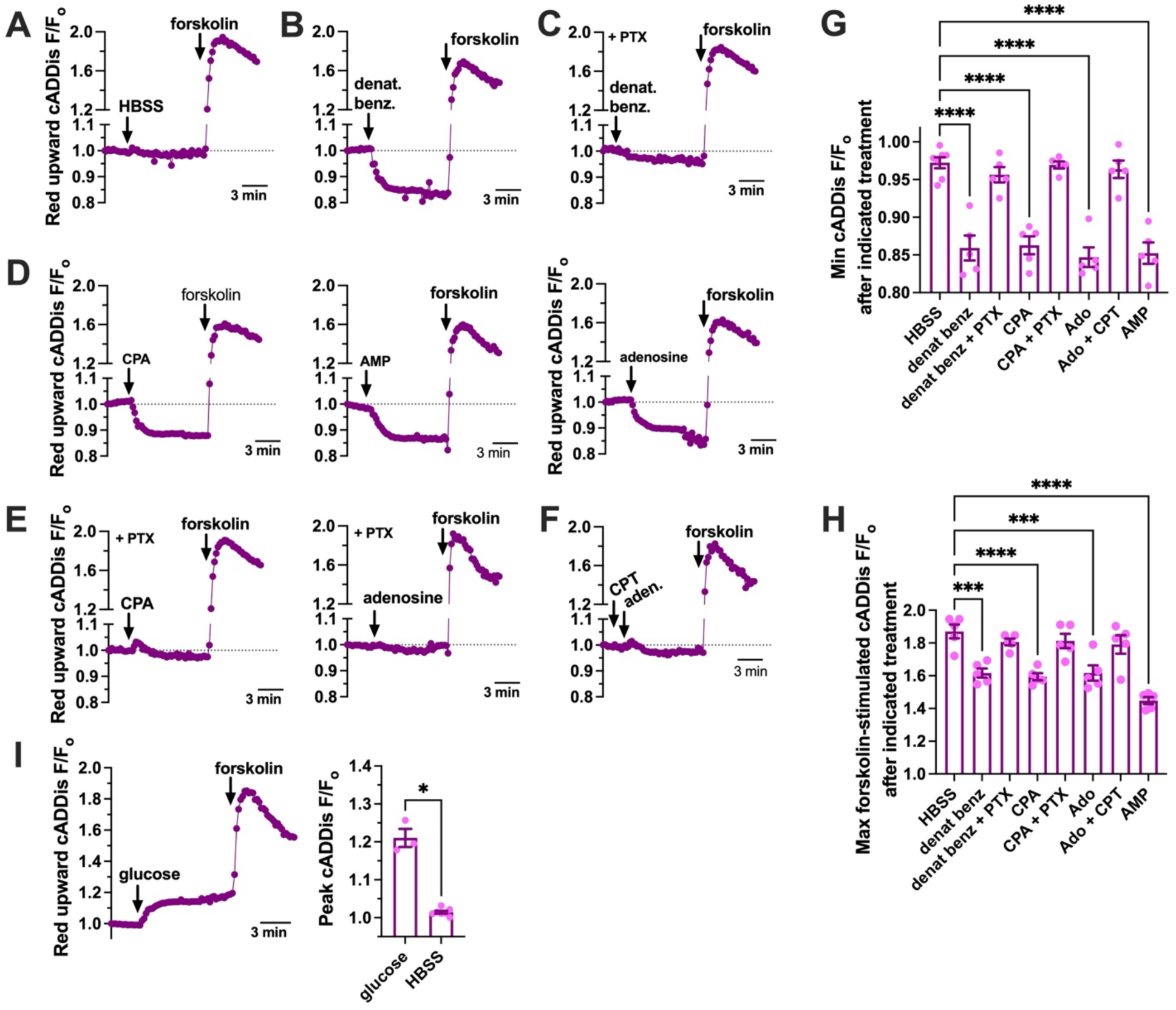
GPCR-induced cAMP changes in Trpm5-GFP-positive mouse nasal tuft cells. **(A-F)** Representative traces showing changes in cADDis fluorescence (cAMP signaling) with the stimuli and inhibitors indicated in the text. **(G-H)** Quantification of the peak decrease in cAMP after 5 min of GPCR stimulation (G) and of the peak increase with forskolin (H). Significance by one-way ANOVA with Bonferroni post test with paired comparisons; \*\*\**p* < 0.001, \*\*\*\**p* < 0.0001; n = 5-6 experiments per condition. **(I)** Traces and summary bar graph showing the increase in cADDis fluorescence (cAMP) with glucose. Significance in the bar graph by Student’s *t* test; \**p* < 0.05; n = 3 independent experiments per condition.

### 3.4 Nasal tuft cells are enriched for *Pde2a*, and PDE2A inhibition raises cAMP and suppresses Ca^2+^ responses

The data above show that Ca^2+^-activating GPCRs in mouse nasal tuft cells also lower cAMP, while T1R2/3 activation raises it. Our prior data suggest that T1R2/3 activation inhibits tuft cell Ca^2+^ responses, which we hypothesized occurred through cAMP. To test this more directly, and to ask whether the cAMP pathway can be used to inhibit tuft cell Ca^2+^ signaling, we examined PDE gene expression in airway tuft cells using publicly available single-cell datasets.^55, 81^ Mouse tracheal tuft cells appeared to be enriched for *Pde2a* (**Figure 5A-C, Supplemental Figures 5 and 6**), agreeing with a previously published study showing Pde2a expression in tuft cells.^21^ The non-selective PDE inhibitor isobutylmethylxanthine (IBMX) and the PDE2A-selective inhibitors PF-05180999^82^ (PF) and BAY 60-7550^83^ (BAY) each increased cAMP in mouse nasal tuft cells (**Figure 5D**). PF and BAY reduced Ca^2+^ responses to denatonium benzoate and CPA, and PF reduced succinate-evoked Ca^2+^ responses (**Figure 5E-F**).

**Figure 5.**
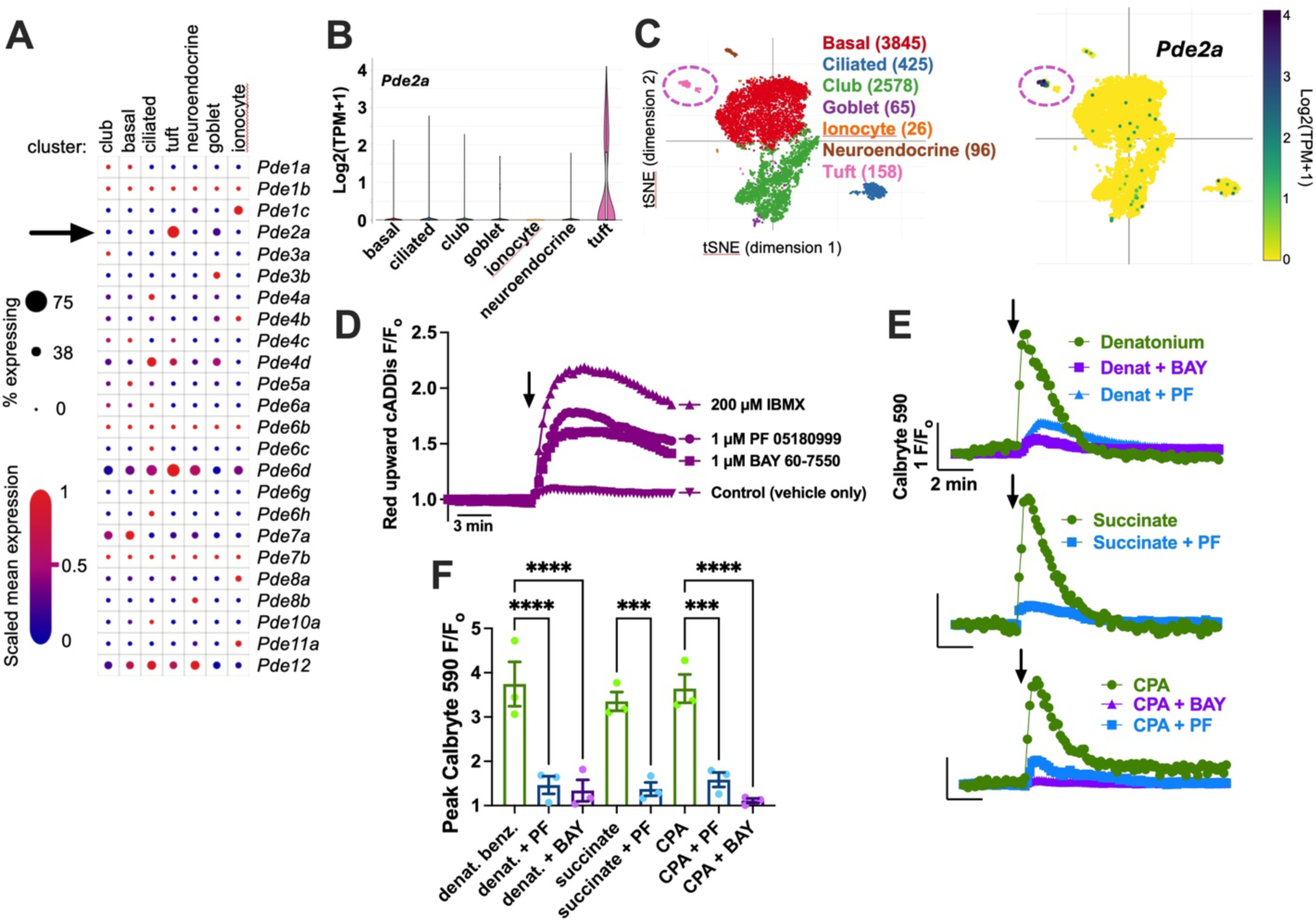
PDE2A inhibition modifies cAMP and Ca^2+^ signaling in mouse nasal Trpm5-GFP-positive tuft cells. **(A-C)** Plot of all *Pde* isoforms (A), *Pde2a* distribution (B), and tSNE plot of *Pde2a* expression (C) using data from Montoro et al.,^55^ showing enrichment of *Pde2a* in mouse tracheal tuft cells. **(D)** Traces of the cADDis response in Trpm5-GFP-positive cells showing an increase in fluorescence (increase in cAMP) with PDE inhibitors. **(E)** Traces of Calbryte 590 showing inhibition of denatonium-, succinate-, and CPA-induced Ca^2+^ signaling by PDE2A-selective inhibitors in Trpm5-GFP-positive cells. **(F)** Summary bar graph of peak Ca^2+^ responses from experiments as in E. Significance by one-way ANOVA with Bonferroni post test with paired comparisons; \*\*\**p* < 0.001, \*\*\*\**p* < 0.0001. All experimental data shown are representative of at least three independent experiments using cells from multiple Trpm5-GFP mice on different days. Data for A-C were accessed and plotted through the Broad Institute Single Cell Portal (https://singlecell.broadinstitute.org/single_cell/study/SCP163/airway-epithelium).

**Figure 6.**
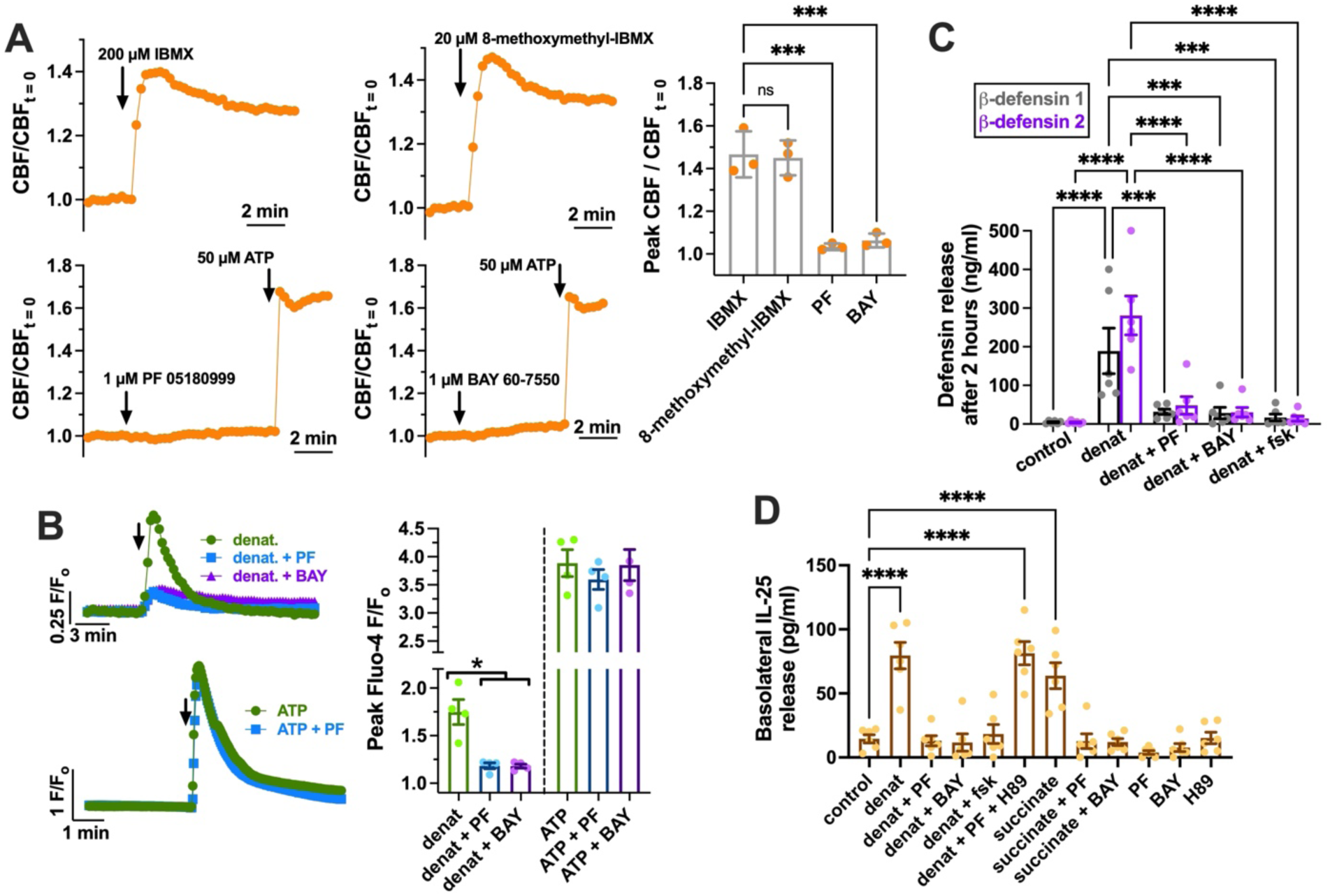
Reduction of human nasal tuft cell GPCR responses by PDE2A-selective inhibitors. **(A)** Representative traces of ciliary beat frequency showing the lack of an increase with PDE2A-selective inhibitors, with quantification in the summary bar graph at right. n = 3 experiments per condition using separate ALI cultures from 3 separate patients with CRSwNP. **(B)** Representative Ca^2+^ responses in human nasal ALIs to the T2R agonist denatonium benzoate (10 mM) or the purinergic agonist ATP (100 µM) as a positive control confirming epithelial responsiveness. The bar graph shows quantification of responses from 4 independent experiments per condition using separate ALIs derived from 4 individual patients with CRSwNP. **(C)** Defensin release quantified by ELISA in ALIs (6 independent experiments per condition using separate ALIs derived from 6 individual patients with CRSwNP) in response to denatonium benzoate, as previously described.^42^ PDE2A-selective inhibitors and forskolin both inhibited defensin release. **(D)** IL-25 release quantified by ELISA as described in the Supporting Information (6 independent experiments per condition using separate ALIs derived from 6 individual patients with CRSwNP, all pre-treated with IL-13 to increase tuft cell numbers, as in Figure 2C-D) in response to denatonium benzoate or succinate. PDE2A inhibition reduced IL-25 release. Significance in A, B, and C by one-way ANOVA with Dunnett’s post test comparing all values to IBMX (A), denatonium (B), or vehicle alone (C). Significance in D by one-way ANOVA with Bonferroni post test. \**p* < 0.05, \*\*\**p* < 0.001, \*\*\*\**p* < 0.0001.

### 3.5 PDE2A inhibition reduces denatonium-evoked Ca^2+^, defensin, and IL-25 responses in human nasal ALI cultures

Single-cell RNAseq Data from the Human Protein Atlas suggests that, while PDE2A is not the most abundant PDE in human tuft cells, it is more highly expressed in human tuft cells than in respiratory ciliated cells, secretory cells, or ionocytes (**Supplemental Figure 7**). To test if PDE2A inhibition might influence human tuft cell response, we first examined the selectivity of PDE2A inhibitors by testing if they increase ciliary beat frequency (CBF) in human nasal epithelial ALI cultures from CRSwNP patients, since cAMP/PKA signaling is well known to increase airway epithelial CBF.^84^ CBF was increased by both non-selective PDE inhibitor IBMX and PDE1A inhibitor 8-methoxymethyl-IBMX (**Figure 6A**), which was previously shown to increase CBF.^85^ However, we saw no increase in CBF with PDE2A inhibitors PF or BAY at the concentrations shown to inhibit mouse nasal tuft cell Ca^2+^ signaling (**Figure 6A**). Both PF and BAY reduced Ca^2+^ responses to denatonium benzoate in human nasal ALIs (**Figure 6B**), which we previously showed involve nasal tuft cells.^42^ Downstream defensin release in response to denatonium benzoate was reduced by PF and BAY, as well as by non-selective elevation of cAMP with forskolin (**Figure 6C**). Denatonium-induced IL-25 release was likewise reduced by PF, BAY, and forskolin, and the effect of PF was reversed by the PKA inhibitor H89 (**Figure 6D**), suggesting that the block of denatonium responses by PF involves PKA. Succinate-induced IL-25 release was also reduced by PF and BAY (**Figure 6D**). Together, these data suggest that targeting the cAMP pathway may be a way to reduce responses from nasal tuft cells by altering Ca^2+^ signaling.

## 4 Discussion

Here we show that T2Rs and other GPCRs that modify tuft cell Ca^2+^ responses also reduce cAMP, likely due to receptor coupling to Gα-gustducin or Gαi pathways. While cAMP levels at baseline are generally assumed to be low, we have also observed in airway epithelial cell lines^40^ and primary human macrophages^86^ that T2Rs can lower baseline cAMP levels. Our data suggest that directly targeting the cAMP pathway could be a way to reduce tuft cell Ca^2+^ signaling and inhibit IL-25 secretion in certain disease settings, including possibly CRSwNP. Finding tuft cell-specific cAMP regulators like PDE isoforms or Gαs-coupled receptors might allow topical therapies to modify cAMP within tuft cells but not surrounding epithelial cells. We are not suggesting from our data that PDE2A is necessarily a good therapeutic target. That conclusion would require further testing both in vitro and in vivo. Our ALI culture system precludes us from understanding how PDE2A inhibitors would affect, for example, endothelial cells, nasal gland cells, and immune cells. While our examination of publicly-available datasets suggests PDE2A is expressed in tuft cells, it is not likely exclusive to tuft cells. However, as a proof-of-principle, our preclinical work here demonstrates potential feasibility of specifically targeting the cAMP pathway in this cell type to support further investigation.

While our data suggest that we could target PDE2 to acutely modify tuft cell GPCR Ca^2+^ signaling, it is unknown if PDE2 activation is part of the tuft cell/GPCR/gustducin pathway. If yes, it remains to be determined how PDE2 is regulated. PDE2 hydrolyzes both cAMP and cGMP and is allosterically activated by cGMP. ^87^ PDE2 is highly expressed in non-taste tissues like the brain^88^ and heart,^87^ and PDE2 modulators have been investigated in mouse models of aging, cognitive decline, brain injury, and heart failure.^89–93^ Gustducin itself is one candidate regulator. It is the third isoform of the transducin Gα family. Transducins typically regulate PDE6 in retinal rod and cone cells to modulate cGMP levels.^94^ Gustducin and transducins may have different PDE specificity, as expression of transducin 1 in mouse taste cells only partially rescues knockout of gustducin,^95^ despite the fact that taste receptors can likely activate GTPase activity of both alpha-gustducin and alpha-transducin similarly.^96, 97^ This suggests that gustducin and transducins may have different downstream effectors. The relevant isoform may also vary by cell type, as tongue taste cells have been suggested to be enriched in PDE3.^98^ Regardless of whether gustducin acts on PDE2, the enzyme can be regulated by other mechanisms, including cGMP binding, Ca^2+^, and protein kinase C.^94, 99, 100^ PDE2 has a hydrophobic N-terminal portion^100^ that can anchor it to lipid rafts^101^ and may help it localize closely with other raft-associated proteins such as adenylyl cyclase and endothelial nitric oxide synthase.^102^ We speculate that this could also help PDE2 localize closely to taste receptors in airway tuft cells. Tuft cells in the intestine are enriched in apically-localized dynamin-2,^103^ which is often associated with lipid rafts.^104–106^ Cholesterol, which is enriched in lipid rafts,^107^ is important for bitter taste receptor function.^108–111^ Lipid rafts are thought to be important for recruitment and trafficking of taste receptors to the taste cell apical membrane on the tongue.^112^ Based on this circumstantial evidence and the expression of PDE2 in airway tuft cells, it is tempting to speculate that PDE2 may be part of the tuft cell GPCR/gustducin pathway, but this requires future experimental investigation.

One practical question for any PDE-directed strategy in the nose is that PDE5 inhibitors such as sildenafil used for erectile dysfunction can contribute to nasal congestion, which is usually attributed to vasodilation.^113^ The off-target activity of these drugs is directed at PDE1, PDE6, and PDE11 rather than PDE2.^114^ Furthermore, the PDE4 inhibitor roflumilast improved nasal airflow and subjective nasal obstruction in a randomized, placebo-controlled crossover study in allergic rhinitis,^115^ and inhibitors of PDE3, PDE4, and PDE5 increased ciliary beat frequency in rabbit maxillary sinus and tracheal explants,^116^ indicating that epithelial and vascular responses to PDE inhibition can differ. We saw no increase in ciliary beat frequency with the PDE2A-selective compounds used here. We therefore have no specific reason to expect congestion from a PDE2A-directed approach, though this has not been tested directly.

Several limitations should be noted. Our conclusions about PDE2A rest on two pharmacological inhibitors, PF-05180999 and BAY 60-7550, without genetic confirmation, so we cannot exclude off-target activity. The absence of a CBF increase with PF and BAY argues against broad cAMP elevation across the epithelium, but it is a negative result and does not by itself establish selectivity for tuft cells. All human ALI cultures were derived from patients with CRSwNP, which is the population of interest but leaves us without a non-polyp comparison. Continued studies are needed to address these and other gaps in our knowledge of how cAMP regulates Ca^2+^ signals and other responses in nasal tuft cells.

## 5 Conclusion

Nasal tuft cell GPCRs that raise Ca^2+^ also lower cAMP, and raising cAMP through PDE2A inhibition reduces tuft cell Ca^2+^ signaling, defensin release, and IL-25 release in vitro. These findings identify the cAMP pathway as a point of control over tuft cell activation and support further in vitro and in vivo investigation of cAMP and Ca^2+^ signaling in tuft cell-driven airway inflammation.

## Acknowledgements

We thank M. Victoria for excellent technical assistance. We thank R. Margolskee (Monell Chemical Senses Center) for originally providing Trpm5-GFP mice and for helpful discussions. We thank K. Foskett (University of Pennsylvania) for allowing us to use the nasal tissue from Trpm5-GFP mice being used for other unrelated experiments.

## References

1. Naclerio R, Baroody F, Bachert C, et al. Clinical Research Needs for the Management of Chronic Rhinosinusitis with Nasal Polyps in the New Era of Biologics: A National Institute of Allergy and Infectious Diseases Workshop. J Allergy Clin Immunol Pract 2020; 8:1532–1549 e1531.

2. Barnes PJ. Targeting cytokines to treat asthma and chronic obstructive pulmonary disease. Nat Rev Immunol 2018; 18:454–466.

3. Laidlaw TM, and Buchheit KM. Biologics in chronic rhinosinusitis with nasal polyposis. Ann Allergy Asthma Immunol 2020; 124:326–332.

4. Holgate ST. Epithelium dysfunction in asthma. J Allergy Clin Immunol 2007; 120:1233–1244; quiz 1245-1236.

5. Stevens WW, Lee RJ, Schleimer RP, et al. Chronic rhinosinusitis pathogenesis. J Allergy Clin Immunol 2015; 136:1442–1453.

6. Roan F, Obata-Ninomiya K, and Ziegler SF. Epithelial cell-derived cytokines: more than just signaling the alarm. J Clin Invest 2019; 129:1441–1451.

7. Mitchell PD, and O’Byrne PM. Biologics and the lung: TSLP and other epithelial cell-derived cytokines in asthma. Pharmacol Ther 2017; 169:104–112.

8. Lee RJ, and Cohen NA. Innate immune function in chronic rhinosinusitis. J Allergy Clin Immunol 2025.

9. Strine MS, and Wilen CB. Tuft cells are key mediators of interkingdom interactions at mucosal barrier surfaces. PLoS Pathog 2022; 18:e1010318.

10. Billipp TE, Nadjsombati MS, and von Moltke J. Tuning tuft cells: new ligands and effector functions reveal tissue-specific function. Curr Opin Immunol 2021; 68:98–106.

11. von Moltke J, Ji M, Liang HE, et al. Tuft-cell-derived IL-25 regulates an intestinal ILC2-epithelial response circuit. Nature 2016; 529:221–225.

12. Howitt MR, Lavoie S, Michaud M, et al. Tuft cells, taste-chemosensory cells, orchestrate parasite type 2 immunity in the gut. Science 2016; 351:1329–1333.

13. Gerbe F, Sidot E, Smyth DJ, et al. Intestinal epithelial tuft cells initiate type 2 mucosal immunity to helminth parasites. Nature 2016; 529:226–230.

14. Krasteva G, Canning BJ, Hartmann P, et al. Cholinergic chemosensory cells in the trachea regulate breathing. Proc Natl Acad Sci U S A 2011; 108:9478–9483.

15. Tizzano M, Gulbransen BD, Vandenbeuch A, et al. Nasal chemosensory cells use bitter taste signaling to detect irritants and bacterial signals. Proc Natl Acad Sci U S A 2010; 107:3210–3215.

16. Saunders CJ, Christensen M, Finger TE, et al. Cholinergic neurotransmission links solitary chemosensory cells to nasal inflammation. Proc Natl Acad Sci U S A 2014; 111:6075–6080.

17. Perniss A, Liu S, Boonen B, et al. Chemosensory Cell-Derived Acetylcholine Drives Tracheal Mucociliary Clearance in Response to Virulence-Associated Formyl Peptides. Immunity 2020; 52:683–699 e611.

18. Hollenhorst MI, Jurastow I, Nandigama R, et al. Tracheal brush cells release acetylcholine in response to bitter tastants for paracrine and autocrine signaling. FASEB J 2020; 34:316–332.

19. Bankova LG, Dwyer DF, Yoshimoto E, et al. The cysteinyl leukotriene 3 receptor regulates expansion of IL-25-producing airway brush cells leading to type 2 inflammation. Sci Immunol 2018; 3.

20. Ualiyeva S, Hallen N, Kanaoka Y, et al. Airway brush cells generate cysteinyl leukotrienes through the ATP sensor P2Y2. Sci Immunol 2020; 5.

21. Ualiyeva S, Lemire E, Aviles EC, et al. Tuft cell-produced cysteinyl leukotrienes and IL-25 synergistically initiate lung type 2 inflammation. Sci Immunol 2021; 6:eabj0474.

22. Baxter BD, Larson ED, Merle L, et al. Transcriptional profiling reveals potential involvement of microvillous TRPM5-expressing cells in viral infection of the olfactory epithelium. BMC Genomics 2021; 22:224.

23. Sell EA, Ortiz-Carpena JF, Herbert DR, et al. Tuft cells in the pathogenesis of chronic rhinosinusitis with nasal polyps and asthma. Ann Allergy Asthma Immunol 2021; 126:143–151.

24. Rane CK, Jackson SR, Pastore CF, et al. Development of solitary chemosensory cells in the distal lung after severe influenza injury. Am J Physiol Lung Cell Mol Physiol 2019; 316:L1141–L1149.

25. Gordon RE, and Kattan M. Absence of cilia and basal bodies with predominance of brush cells in the respiratory mucosa from a patient with immotile cilia syndrome. Ultrastruct Pathol 1984; 6:45–49.

26. Cerezo L, and Price G. Absence of cilia and basal bodies with predominance of brush cells in the respiratory mucosa from a patient with immotile cilia syndrome. Ultrastruct Pathol 1985; 8:381–382.

27. DiMaio MF, Dische R, Gordon RE, et al. Alveolar brush cells in an infant with desquamative interstitial pneumonitis. Pediatr Pulmonol 1988; 4:185–191.

28. Meyrick B, and Reid L. The alveolar brush cell in rat lung--a third pneumonocyte. J Ultrastruct Res 1968; 23:71–80.

29. Reid L, Meyrick B, Antony VB, et al. The mysterious pulmonary brush cell: a cell in search of a function. Am J Respir Crit Care Med 2005; 172:136–139.

30. Patel NN, Triantafillou V, Maina IW, et al. Fungal extracts stimulate solitary chemosensory cell expansion in noninvasive fungal rhinosinusitis. Int Forum Allergy Rhinol 2019; 9:730–737.

31. Patel NN, Kohanski MA, Maina IW, et al. Solitary chemosensory cells producing interleukin-25 and group-2 innate lymphoid cells are enriched in chronic rhinosinusitis with nasal polyps. Int Forum Allergy Rhinol 2018.

32. Kohanski MA, Workman AD, Patel NN, et al. Solitary chemosensory cells are a primary epithelial source of IL-25 in patients with chronic rhinosinusitis with nasal polyps. J Allergy Clin Immunol 2018.

33. Shin HW, Kim DK, Park MH, et al. IL-25 as a novel therapeutic target in nasal polyps of patients with chronic rhinosinusitis. J Allergy Clin Immunol 2015; 135:1476–1485 e1477.

34. Finger TE, Bottger B, Hansen A, et al. Solitary chemoreceptor cells in the nasal cavity serve as sentinels of respiration. Proc Natl Acad Sci U S A 2003; 100:8981–8986.

35. Tizzano M, Cristofoletti M, Sbarbati A, et al. Expression of taste receptors in solitary chemosensory cells of rodent airways. BMC Pulm Med 2011; 11:3.

36. Gulbransen BD, Clapp TR, Finger TE, et al. Nasal solitary chemoreceptor cell responses to bitter and trigeminal stimulants in vitro. J Neurophysiol 2008; 99:2929–2937.

37. Lee RJ, and Cohen NA. Taste receptors in innate immunity. Cell Mol Life Sci 2015; 72:217–236.

38. Jaggupilli A, Singh N, Jesus VC, et al. Characterization of the Binding Sites for Bacterial Acyl Homoserine Lactones (AHLs) on Human Bitter Taste Receptors (T2Rs). ACS Infect Dis 2018; 4:1146–1156.

39. Medapati MR, Singh N, Bhagirath AY, et al. Bitter taste receptor T2R14 detects quorum sensing molecules from cariogenic Streptococcus mutans and mediates innate immune responses in gingival epithelial cells. FASEB J 2021; 35:e21375.

40. Freund JR, Mansfield CJ, Doghramji LJ, et al. Activation of airway epithelial bitter taste receptors by Pseudomonas aeruginosa quinolones modulates calcium, cyclic-AMP, and nitric oxide signaling. J Biol Chem 2018; 293:9824–9840.

41. Lin W, Ogura T, Margolskee RF, et al. TRPM5-expressing solitary chemosensory cells respond to odorous irritants. J Neurophysiol 2008; 99:1451–1460.

42. Lee RJ, Kofonow JM, Rosen PL, et al. Bitter and sweet taste receptors regulate human upper respiratory innate immunity. J Clin Invest 2014; 124:1393–1405.

43. Lee RJ, Hariri BM, McMahon DB, et al. Bacterial d-amino acids suppress sinonasal innate immunity through sweet taste receptors in solitary chemosensory cells. Sci Signal 2017; 10:eaam7703.

44. Wong GT, Gannon KS, and Margolskee RF. Transduction of bitter and sweet taste by gustducin. Nature 1996; 381:796–800.

45. Ming D, Ninomiya Y, and Margolskee RF. Blocking taste receptor activation of gustducin inhibits gustatory responses to bitter compounds. Proc Natl Acad Sci U S A 1999; 96:9903–9908.

46. Ruiz-Avila L, Wong GT, Damak S, et al. Dominant loss of responsiveness to sweet and bitter compounds caused by a single mutation in alpha -gustducin. Proc Natl Acad Sci U S A 2001; 98:8868–8873.

47. Masubuchi Y, Nakagawa Y, Ma J, et al. A novel regulatory function of sweet taste-sensing receptor in adipogenic differentiation of 3T3-L1 cells. PLoS One 2013; 8:e54500.

48. Naim M, Ronen T, Striem BJ, et al. Adenylate cyclase responses to sucrose stimulation in membranes of pig circumvallate taste papillae. Comp Biochem Physiol B 1991; 100:455–458.

49. Dyer J, Salmon KS, Zibrik L, et al. Expression of sweet taste receptors of the T1R family in the intestinal tract and enteroendocrine cells. Biochem Soc Trans 2005; 33:302–305.

50. Trubey KR, Culpepper S, Maruyama Y, et al. Tastants evoke cAMP signal in taste buds that is independent of calcium signaling. Am J Physiol Cell Physiol 2006; 291:C237–244.

51. Nakagawa Y, Nagasawa M, Yamada S, et al. Sweet taste receptor expressed in pancreatic beta-cells activates the calcium and cyclic AMP signaling systems and stimulates insulin secretion. PLoS One 2009; 4:e5106.

52. Barham HP, Cooper SE, Anderson CB, et al. Solitary chemosensory cells and bitter taste receptor signaling in human sinonasal mucosa. Int Forum Allergy Rhinol 2013; 3:450–457.

53. Howitt MR, Cao YG, Gologorsky MB, et al. The Taste Receptor TAS1R3 Regulates Small Intestinal Tuft Cell Homeostasis. Immunohorizons 2020; 4:23–32.

54. Sell EA, Tan LH, Lin C, et al. Microbial metabolite succinate activates solitary chemosensory cells in the human sinonasal epithelium. Int Forum Allergy Rhinol 2022.

55. Montoro DT, Haber AL, Biton M, et al. A revised airway epithelial hierarchy includes CFTR-expressing ionocytes. Nature 2018; 560:319–324.

56. Huang L, Shanker YG, Dubauskaite J, et al. Ggamma13 colocalizes with gustducin in taste receptor cells and mediates IP3 responses to bitter denatonium. Nat Neurosci 1999; 2:1055–1062.

57. Clapp TR, Stone LM, Margolskee RF, et al. Immunocytochemical evidence for co-expression of Type III IP3 receptor with signaling components of bitter taste transduction. BMC Neurosci 2001; 2:6.

58. Yan W, Sunavala G, Rosenzweig S, et al. Bitter taste transduced by PLC-beta(2)-dependent rise in IP(3) and alpha-gustducin-dependent fall in cyclic nucleotides. Am J Physiol Cell Physiol 2001; 280:C742–751.

59. Clapp TR, Trubey KR, Vandenbeuch A, et al. Tonic activity of Galpha-gustducin regulates taste cell responsivity. FEBS Lett 2008; 582:3783–3787.

60. Liu M, and Simon MI. Regulation by cAMP-dependent protein kinease of a G-protein-mediated phospholipase C. Nature 1996; 382:83–87.

61. Giovannucci DR, Groblewski GE, Sneyd J, et al. Targeted phosphorylation of inositol 1,4,5-trisphosphate receptors selectively inhibits localized Ca2+ release and shapes oscillatory Ca2+ signals. J Biol Chem 2000; 275:33704–33711.

62. Soulsby MD, and Wojcikiewicz RJ. The type III inositol 1,4,5-trisphosphate receptor is phosphorylated by cAMP-dependent protein kinase at three sites. Biochem J 2005; 392:493–497.

63. Dyer JL, Mobasheri H, Lea EJ, et al. Differential effect of PKA on the Ca2+ release kinetics of the type I and III InsP3 receptors. Biochem Biophys Res Commun 2003; 302:121–126.

64. Clapp TR, Medler KF, Damak S, et al. Mouse taste cells with G protein-coupled taste receptors lack voltage-gated calcium channels and SNAP-25. BMC Biol 2006; 4:7.

65. Lee RJ, Chen B, Redding KM, et al. Mouse nasal epithelial innate immune responses to Pseudomonas aeruginosa quorum-sensing molecules require taste signaling components. Innate Immun 2014; 20:606–617.

66. Kuek LE, McMahon DB, Ma RZ, et al. Cilia Stimulatory and Antibacterial Activities of T2R Bitter Taste Receptor Agonist Diphenhydramine: Insights into Repurposing Bitter Drugs for Nasal Infections. Pharmaceuticals (Basel) 2022; 15.

67. Kouakou YI, Thompson JC, Tan LH, et al. Hops bitter beta-acids have antibacterial effects against sinonasal Staphylococcus aureus but also induce sinonasal cilia and mitochondrial dysfunction. Int Forum Allergy Rhinol 2024; 15:287–302.

68. Thompson JC, Park A, Kouakou YI, et al. Mechanisms of Mitochondrial Toxicity and Cytotoxicity Caused by Pseudomonas aeruginosa Pyocyanin in Human Nasal Epithelial Cells. Int Forum Allergy Rhinol 2026; 16:451–466.

69. Sisson JH, Stoner JA, Ammons BA, et al. All-digital image capture and whole-field analysis of ciliary beat frequency. J Microsc 2003; 211:103–111.

70. Perez CA, Huang L, Rong M, et al. A transient receptor potential channel expressed in taste receptor cells. Nat Neurosci 2002; 5:1169–1176.

71. Lee RJ, and Cohen NA. Sinonasal solitary chemosensory cells “taste” the upper respiratory environment to regulate innate immunity. Am J Rhinol Allergy 2014; 28:366–373.

72. Hollenhorst MI, Nandigama R, Evers SB, et al. Bitter taste signaling in tracheal epithelial brush cells elicits innate immune responses to bacterial infection. J Clin Invest 2022.

73. Kotas ME, Moore CM, Gurrola JG, 2nd, et al. IL-13-programmed airway tuft cells produce PGE2, which promotes CFTR-dependent mucociliary function. JCI Insight 2022; 7.

74. Deng J, Tan LH, Kohanski MA, et al. Solitary chemosensory cells are innervated by trigeminal nerve endings and autoregulated by cholinergic receptors. Int Forum Allergy Rhinol 2021; 11:877–884.

75. Yamashita J, Ohmoto M, Yamaguchi T, et al. Skn-1a/Pou2f3 functions as a master regulator to generate Trpm5-expressing chemosensory cells in mice. PLoS One 2017; 12:e0189340.

76. Ualiyeva S, Lemire E, Wong C, et al. A nasal cell atlas reveals heterogeneity of tuft cells and their role in directing olfactory stem cell proliferation. Sci Immunol 2024; 9:eabq4341.

77. Krasteva-Christ G, Soultanova A, Schutz B, et al. Identification of cholinergic chemosensory cells in mouse tracheal and laryngeal glandular ducts. Int Immunopharmacol 2015; 29:158–165.

78. Krasteva G, Hartmann P, Papadakis T, et al. Cholinergic chemosensory cells in the auditory tube. Histochem Cell Biol 2012; 137:483–497.

79. Roper SD. Encoding Taste: From Receptors to Perception. Handb Exp Pharmacol 2022; 275:53–90.

80. Ozeck M, Brust P, Xu H, et al. Receptors for bitter, sweet and umami taste couple to inhibitory G protein signaling pathways. Eur J Pharmacol 2004; 489:139–149.

81. Plasschaert LW, Zilionis R, Choo-Wing R, et al. A single-cell atlas of the airway epithelium reveals the CFTR-rich pulmonary ionocyte. Nature 2018; 560:377–381.

82. Helal CJ, Arnold E, Boyden T, et al. Identification of a Potent, Highly Selective, and Brain Penetrant Phosphodiesterase 2A Inhibitor Clinical Candidate. J Med Chem 2018; 61:1001–1018.

83. Boess FG, Hendrix M, van der Staay FJ, et al. Inhibition of phosphodiesterase 2 increases neuronal cGMP, synaptic plasticity and memory performance. Neuropharmacology 2004; 47:1081–1092.

84. Kuek LE, and Lee RJ. First contact: the role of respiratory cilia in host-pathogen interactions in the airways. Am J Physiol Lung Cell Mol Physiol 2020; 319:L603–L619.

85. Kogiso H, Hosogi S, Ikeuchi Y, et al. A low [Ca(2+)](i)-induced enhancement of cAMP-activated ciliary beating by PDE1A inhibition in mouse airway cilia. Pflugers Arch 2017; 469:1215–1227.

86. Gopallawa I, Freund JR, and Lee RJ. Bitter taste receptors stimulate phagocytosis in human macrophages through calcium, nitric oxide, and cyclic-GMP signaling. Cell Mol Life Sci 2021; 78:271–286.

87. Sadek MS, Cachorro E, El-Armouche A, et al. Therapeutic Implications for PDE2 and cGMP/cAMP Mediated Crosstalk in Cardiovascular Diseases. Int J Mol Sci 2020; 21.

88. Delhaye S, and Bardoni B. Role of phosphodiesterases in the pathophysiology of neurodevelopmental disorders. Mol Psychiatry 2021; 26:4570–4582.

89. Reneerkens OAH, Rutten K, Bollen E, et al. Inhibition of phoshodiesterase type 2 or type 10 reverses object memory deficits induced by scopolamine or MK-801. Behav Brain Res 2013; 236:16–22.

90. Domek-Lopacinska K, and Strosznajder JB. The effect of selective inhibition of cyclic GMP hydrolyzing phosphodiesterases 2 and 5 on learning and memory processes and nitric oxide synthase activity in brain during aging. Brain Res 2008; 1216:68–77.

91. Bechay KR, Abduljawad N, Latifi S, et al. PDE2A Inhibition Enhances Axonal Sprouting, Functional Connectivity, and Recovery after Stroke. J Neurosci 2022; 42:8225–8236.

92. Vettel C, Lindner M, Dewenter M, et al. Phosphodiesterase 2 Protects Against Catecholamine-Induced Arrhythmia and Preserves Contractile Function After Myocardial Infarction. Circ Res 2017; 120:120–132.

93. Mehel H, Emons J, Vettel C, et al. Phosphodiesterase-2 is up-regulated in human failing hearts and blunts beta-adrenergic responses in cardiomyocytes. J Am Coll Cardiol 2013; 62:1596–1606.

94. Omori K, and Kotera J. Overview of PDEs and their regulation. Circ Res 2007; 100:309–327.

95. He W, Danilova V, Zou S, et al. Partial rescue of taste responses of alpha-gustducin null mice by transgenic expression of alpha-transducin. Chem Senses 2002; 27:719–727.

96. Hoon MA, Northup JK, Margolskee RF, et al. Functional expression of the taste specific G-protein, alpha-gustducin. Biochem J 1995; 309 (Pt 2):629–636.

97. Ruiz-Avila L, McLaughlin SK, Wildman D, et al. Coupling of bitter receptor to phosphodiesterase through transducin in taste receptor cells. Nature 1995; 376:80–85.

98. McLaughlin SK, McKinnon PJ, Spickofsky N, et al. Molecular cloning of G proteins and phosphodiesterases from rat taste cells. Physiol Behav 1994; 56:1157–1164.

99. Geoffroy V, Fouque F, Nivet V, et al. Activation of a cGMP-stimulated cAMP phosphodiesterase by protein kinase C in a liver Golgi-endosomal fraction. Eur J Biochem 1999; 259:892–900.

100. Yang Q, Paskind M, Bolger G, et al. A novel cyclic GMP stimulated phosphodiesterase from rat brain. Biochem Biophys Res Commun 1994; 205:1850–1858.

101. Noyama K, and Maekawa S. Localization of cyclic nucleotide phosphodiesterase 2 in the brain-derived Triton-insoluble low-density fraction (raft). Neurosci Res 2003; 45:141–148.

102. Mongillo M, Tocchetti CG, Terrin A, et al. Compartmentalized phosphodiesterase-2 activity blunts beta-adrenergic cardiac inotropy via an NO/cGMP-dependent pathway. Circ Res 2006; 98:226–234.

103. Silverman JB, Krystofiak EE, Caplan LR, et al. Organization of a cytoskeletal superstructure in the apical domain of intestinal tuft cells. J Cell Biol 2024; 223.

104. Vassilieva EV, Gerner-Smidt K, Ivanov AI, et al. Lipid rafts mediate internalization of beta1-integrin in migrating intestinal epithelial cells. Am J Physiol Gastrointest Liver Physiol 2008; 295:G965–976.

105. Wang Y, Zhang Y, Zuo W, et al. Avian Reovirus sigmaB Interacts with Caveolin-1 in Lipid Rafts during Dynamin-Dependent Caveolae-Mediated Endocytosis. Viruses 2022; 14.

106. Liu YW, Lukiyanchuk V, and Schmid SL. Common membrane trafficking defects of disease-associated dynamin 2 mutations. Traffic 2011; 12:1620–1633.

107. Warda M, Tekin S, Gamal M, et al. Lipid rafts: novel therapeutic targets for metabolic, neurodegenerative, oncological, and cardiovascular diseases. Lipids Health Dis 2025; 24:147.

108. Shaik FA, Medapati MR, and Chelikani P. Cholesterol modulates the signaling of chemosensory bitter taste receptor T2R14 in human airway cells. Am J Physiol Lung Cell Mol Physiol 2019; 316:L45–L57.

109. Pydi SP, Jafurulla M, Wai L, et al. Cholesterol modulates bitter taste receptor function. Biochim Biophys Acta 2016; 1858:2081–2087.

110. Hu X, Ao W, Gao M, et al. Bitter taste TAS2R14 activation by intracellular tastants and cholesterol. Nature 2024.

111. Kim Y, Gumpper RH, Liu Y, et al. Bitter taste receptor activation by cholesterol and an intracellular tastant. Nature 2024; 628:664–671.

112. Ilegems E, Iwatsuki K, Kokrashvili Z, et al. REEP2 enhances sweet receptor function by recruitment to lipid rafts. J Neurosci 2010; 30:13774–13783.

113. Moreira SG, Brannigan RE, Spitz A, et al. Side-effect profile of sildenafil citrate (Viagra) in clinical practice. Urology 2000; 56:474–476.

114. Bischoff E. Potency, selectivity, and consequences of nonselectivity of PDE inhibition. Int J Impot Res 2004; 16:S11–S14.

115. Schmidt BMW, Kusma M, Feuring M, et al. The phosphodiesterase 4 inhibitor roflumilast is effective in the treatment of allergic rhinitis. J Allergy Clin Immunol 2001; 108:530–536.

116. Cervin A, Lindgren S. The effect of selective phosphodiesterase inhibitors on mucociliary activity in the upper and lower airways in vitro. Auris Nasus Larynx 1998; 25:269–276.

