## Supplemental Material for "Targeting cyclic-AMP/phosphodiesterase signaling to quiet nasal tuft cell calcium responses"

### **Supplemental Methods**

#### **S1. Reagents and solutions**

Hydrophobic compounds were prepared as stock solutions in DMSO, and vehicle controls were matched to the highest DMSO concentration used in a given experiment. Live cell imaging was carried out in Hank's balanced salt solution (HBSS) containing (in mM) 137 NaCl, 5 KCl, 0.4 KH<sub>2</sub>PO<sub>4</sub>, 0.3 Na<sub>2</sub>HPO<sub>4</sub>, 5.6 glucose, 1.8 CaCl<sub>2</sub>, 1.5 MgCl<sub>2</sub>, and 20 HEPES, adjusted to pH 7.4. PF-05180999 (Cat # 30374), BAY 60-7550 (Cat # 10011135), isobutylmethylxanthine (IBMX; Cat # 13347), 8-Methoxymethyl-IBMX (8-methoxymethyl-3-isobutyl-1-methylxanthine; Cat #39151), U73122 (Cat # 70740), U73343 (Cat #17339), pertussis toxin (Cat # 23221), H89 (Cat # 10010556), forskolin (Cat # 11018), 8-cyclopentyltheophylline (8-cyclopentyl-1,3-dimethylxanthine; Cat # 29870), and N<sup>6</sup>-cyclopentyladenosine (Cat # 21448) were all from Cayman Chemical (Ann Arbor, MI). Denatonium benzoate (Cat # D5765) and succinate (Cat #S2378) were from MilliporeSigma (Burlington, MA)

Defensins were measured with human β-defensin-1 (ThermoFisher 900-K202K) and β-defensin-2 (ThermoFisher 900-K172K) ELISA development kits (Peprotech/ThermoFisher, Waltham MA), following the protocol we have described previously.<sup>1</sup> For IL-25 ELISA (ab272200, Abcam, Cambridge MA), ALI inserts were laid on the base of a glass-bottom 6-well plate and held down with the hollowed-out cap of a 15 mL conical tube (Corning, Tewksbury, MA). This trapped a small volume of fluid, roughly 50 μL, between the glass and the underside of the insert. About 25 μL of that fluid was recovered and assayed according to the manufacturer's instructions.

#### **S2. Mouse nasal cell culture and imaging**

ALI cultures were established from excised nasal septa as described previously.<sup>2</sup> Epithelial cells were released by digestion with collagenase and pronase and then seeded onto Costar 6.5 mm transwell permeable supports with the apical surface submerged. Cells reached confluence by day 7, at which point apical medium was withdrawn and feeding continued from the basolateral compartment alone. Differentiation and ciliogenesis were evident 10 to 14 days after air exposure, and cultures were used between 28 and 35 days at ALI, except when earlier time points are indicated in Figure 2B. GFP-labeled tuft cells were imaged on an Olympus IX-83 microscope with a standard FITC filter set (Chroma Technologies, Bellows Falls VT), an Orca Flash 4.0 camera (Hamamatsu, Tokyo, Japan), a 10x/0.5 NA or 60x/1.4 NA objective, and MetaFluor software (Molecular Devices, San Jose, CA). Where indicated, mouse nasal ALI cultures were pre-treated basolaterally after differentiation with 10 ng/ml recombinant mouse IL-13 (StemCell Technologies, # 78030) for 10 days before experiments.

For experiments on isolated cells, nasal septa were minced with scissors in $\text{Ca}^{2+}$-containing Tyrode solution (in mM: 140 NaCl, 5 KCl, 2 $\text{CaCl}_2$, 1 $\text{MgCl}_2$, 10 glucose, 5 Na-pyruvate, and 10 HEPES, adjusted to pH 7.4 with NaOH) supplemented with Dispase II (2 mg/mL; Roche, Basel, Switzerland), collagenase P (1 mg/mL; Roche), trypsin inhibitor (1 mg/mL; MilliporeSigma), elastase (0.2 mg/mL; Sigma), and DNase I (10 $\mu\text{g/mL}$; Roche). Tissue was incubated for 45 min with gentle shaking, after which cells were freed by gentle trituration through a glass Pasteur pipet. The supernatant was set aside and the remaining tissue was incubated a further 15 min in $\text{Ca}^{2+}$-free Tyrode solution (in mM: 140 NaCl, 5 KCl, 5 EGTA, 10 glucose, 5 Na-pyruvate, and 10 HEPES, adjusted to pH 7.4 with NaOH) and triturated again. The two supernatants were combined and cells were recovered by gentle centrifugation ($800 \times g$, 2 min).

For $\text{Ca}^{2+}$ imaging, isolated cells were plated on poly-D-ornithine-coated 8-well chamberslides (CellVis, Mountain View, CA), and left to attach for approximately 30 min. They were then loaded with 10 $\mu\text{M}$ Calbryte 590 AM (AAT Bioquest, Pleasanton, CA) for 30 min at room temperature in $\text{Ca}^{2+}$-containing Tyrode solution and washed into glucose-free Tyrode solution (in mM: 140 NaCl, 5 KCl, 2 $\text{CaCl}_2$, 1 $\text{MgCl}_2$, 5 Na-pyruvate, and 10 HEPES, adjusted to pH 7.4 with NaOH) for imaging.

For cAMP imaging with Red Upward cADDIs (Montana Molecular, Bozeman, MT), isolated cells were plated on chamberslides coated with 1:30 Matrigel in 150 $\mu\text{L}$ high-glucose DMEM (Gibco) containing 10% FBS, 1.8 $\mu\text{L}$ sodium butyrate (500 mM stock in $\text{H}_2\text{O}$, supplied with the Red Upward cADDIs kit), and 15 $\mu\text{L}$ cADDIs biosensor in BacMam. Cells were washed into $\text{Ca}^{2+}$-containing Tyrode solution and imaged 18 to 24 h after plating, then washed into glucose-free Tyrode solution immediately before imaging. Both Calbryte 590 and cADDIs were imaged on an Olympus IX-83 microscope with a standard TRITC filter set (Chroma Technologies), an Orca Flash 4.0 camera (Hamamatsu), a 20x/0.8 NA objective, and Metafluor software (Molecular Devices). A FITC filter set was used in the same sessions to identify GFP-positive cells.

#### **S3. Human nasal cell culture and imaging**

Human nasal epithelial cells (HNECs) were released from excised tissue by enzymatic digestion.<sup>3</sup> Tissue was cut into small pieces and held for 1 h at 37 °C in Minimal Essential Medium with Earle's salts and L-glutamine (MEM; Corning, Glendale, AZ, USA) containing 1.4 mg/mL pronase (MilliporeSigma) and 0.1 mg/mL DNase (MilliporeSigma). Digestion was quenched with FBS, released cells were separated from residual tissue through a strainer, and the cells were washed twice in MEM with 10% FBS. Pellets were resuspended in complete PneumaCult-Ex Plus medium (Stemcell Technologies, Vancouver, BC, Canada) prepared per the manufacturer's instructions and supplemented with 1% (v/v) penicillin/streptomycin (Corning). Suspensions were held in T-25 flasks at 37 °C and 5% $\text{CO}_2$ under humidified conditions for 2 h so that non-epithelial cells could adhere and be discarded. Supernatants containing HNECs were moved to fresh T-25 flasks, left overnight, and given fresh medium the next morning. During expansion, cells were fed with PneumaCult-Ex Plus containing 1% (v/v) penicillin/streptomycin, 0.125 µg/mL amphotericin B, 10 µM of the ROCK inhibitor Y27632, and 1 µM DMH-1. Expansion medium was replaced every 2 to 3 days, and cells were passaged once at least 80% confluent.

For ALI culture, HNECs were seeded onto Transwell inserts (Corning; 0.33 cm<sup>2</sup> transparent PTFE membrane, 0.4 µm pore size) and grown submerged until fully confluent. The apical surface was then exposed to air for a minimum of three weeks to permit differentiation. Throughout the ALI phase, cultures were fed basolaterally every 2 to 3 days with complete PneumaCult ALI medium (Stemcell) supplemented with 1% (v/v) penicillin/streptomycin and 0.125 µg/mL amphotericin B. Where indicated, differentiated nasal ALI cultures were pretreated basolaterally with 10 ng/mL recombinant human IL-13 (StemCell Technologies, Cat # 78029) for 10 days before experiments

For Ca<sup>2+</sup> imaging, HNEC ALIs were loaded from the apical side with 10 µM Fluo-4 AM in glucose-free HBSS containing 295 µg/mL probenecid and 394 µg/mL Pluronic F127 for 2 h at room temperature in the dark, as described previously.<sup>1, 4, 5</sup> Ca<sup>2+</sup> was imaged on an Olympus IX-83 microscope (10x/0.40 NA objective) with FITC filters (Chroma Technologies), an Orca Flash 4.0 sCMOS camera (Hamamatsu, Tokyo, Japan), and MetaFluor software (Molecular Devices, Sunnyvale, CA, USA). Ciliary beat frequency was measured at room temperature using Sisson-Ammons Video Analysis software,<sup>6</sup> a Nikon Diaphot microscope (Modulation Optics Inc. HMC 20x LWD 0.4 NA 160/0-2 objective), and a Basler acA1300-200um camera (Ahrensburg, Germany; 120 frames per second) fitted with a 0.7x DXM lens (Nikon, Tokyo, Japan).

### Supplemental Figures

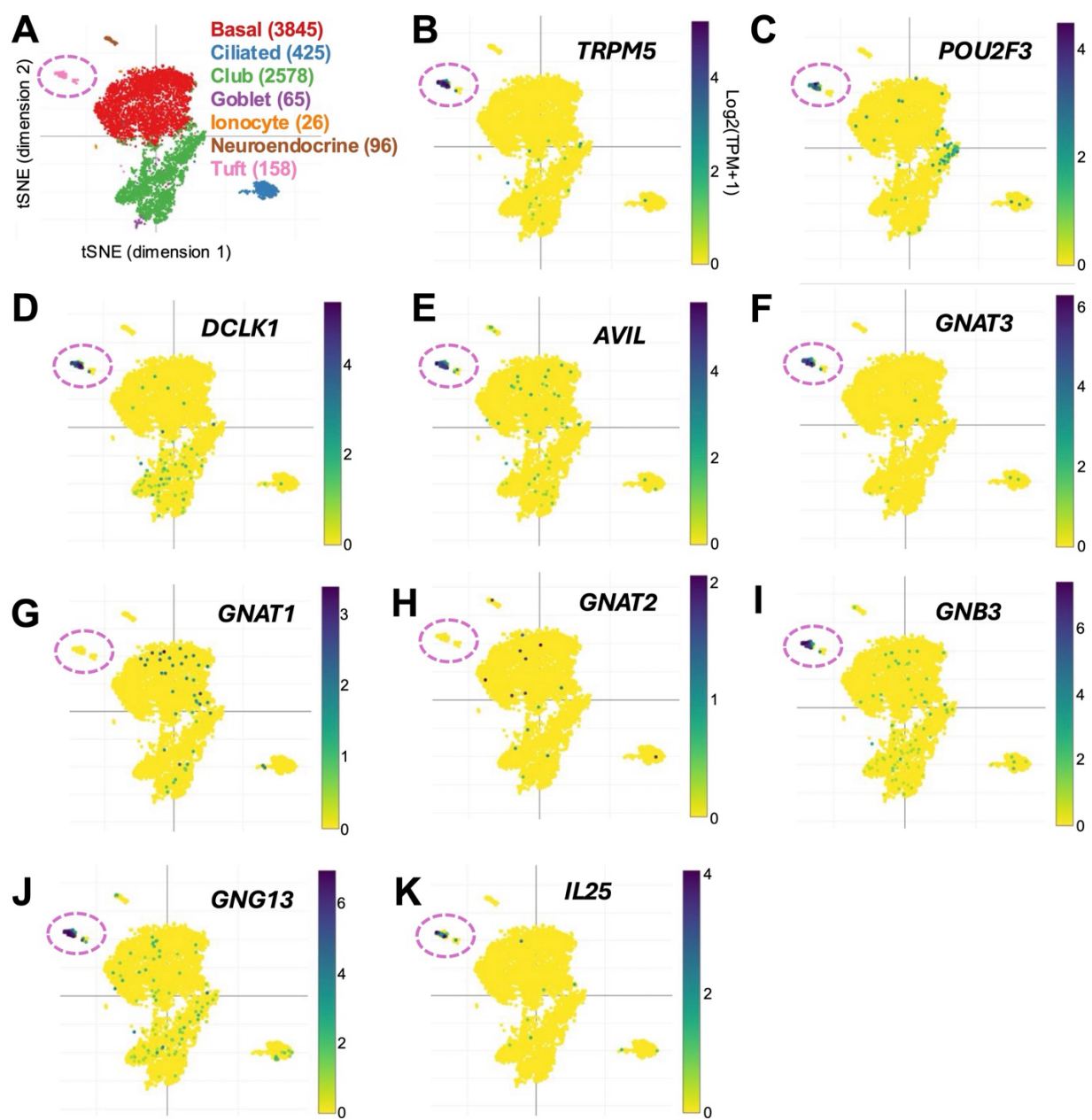

**Supplemental Figure 1.** tSNE plots of gene expression data from Montoro et al.,<sup>7</sup> showing enrichment of tuft cell markers in *Trpm5*-positive mouse tracheal cells (A-B): *Pou2f3* (C), *Dclk1* (D), and *Avil* (E). Tuft cells were enriched for the gene encoding gustducin, *Gnat3* (F), but not for *Gnat1* or *Gnat2* (G-H). The Gβ and Gγ subunit genes *Gnb3* (I) and *Gng13* (J) were also expressed, along with downstream *Il25* (K). Data were accessed and plotted through the Broad Institute Single Cell Portal (https://singlecell.broadinstitute.org/single_cell/study/SCP163/airway-epithelium).

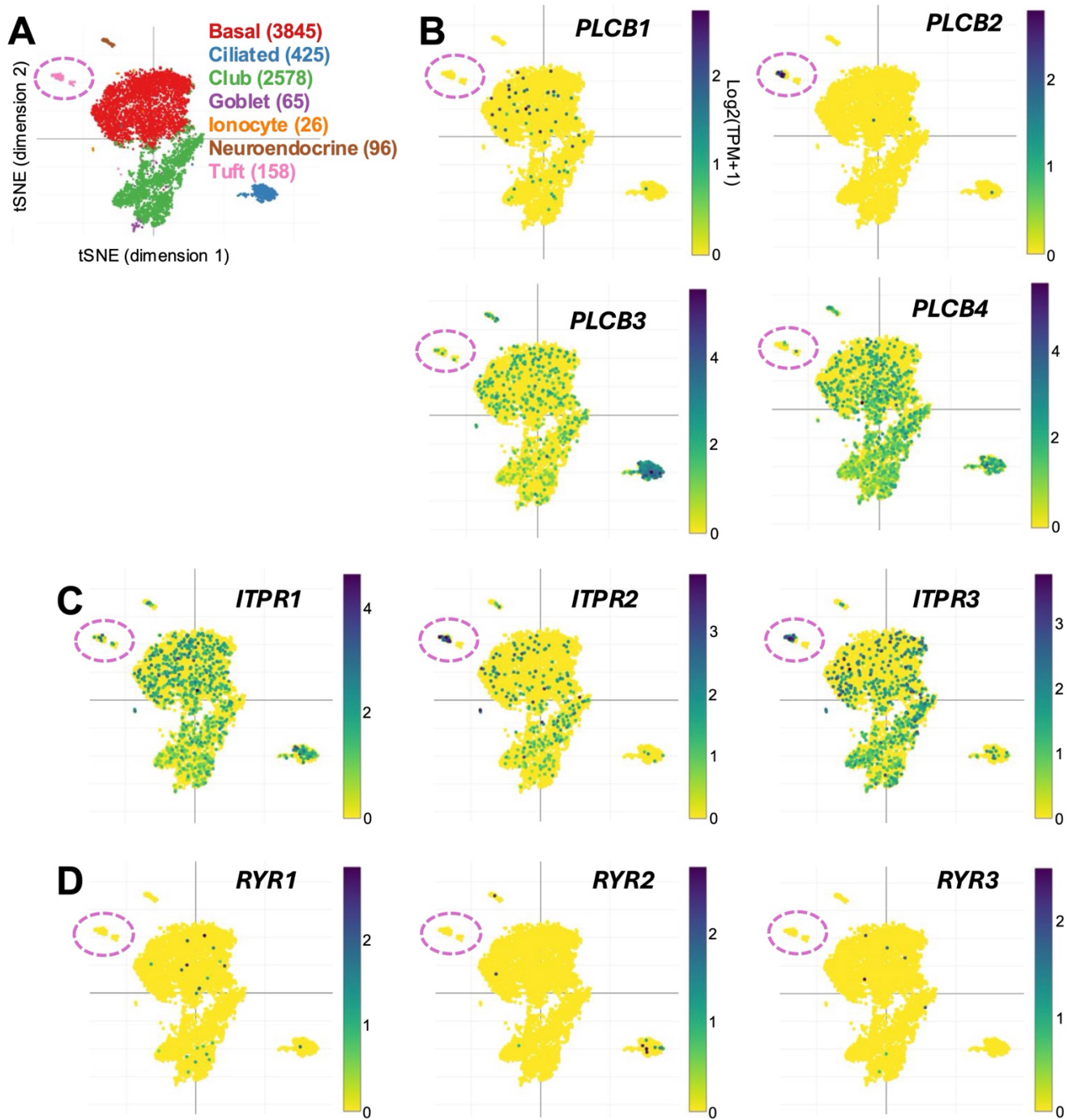

**Supplemental Figure 2.** tSNE plots of gene expression data from Montoro et al.,<sup>7</sup> showing enrichment of taste signaling components in mouse tracheal tuft cells (A), including phospholipase C isoform $\beta 2$ (*Plcb2*; B) and the type III inositol trisphosphate receptor (*Itpr3*; C). Ryanodine receptors, another class of intracellular $\text{Ca}^{2+}$ release channel, were not detected (D). Data were accessed and plotted through the Broad Institute Single Cell Portal (https://singlecell.broadinstitute.org/single_cell/study/SCP163/airway-epithelium).

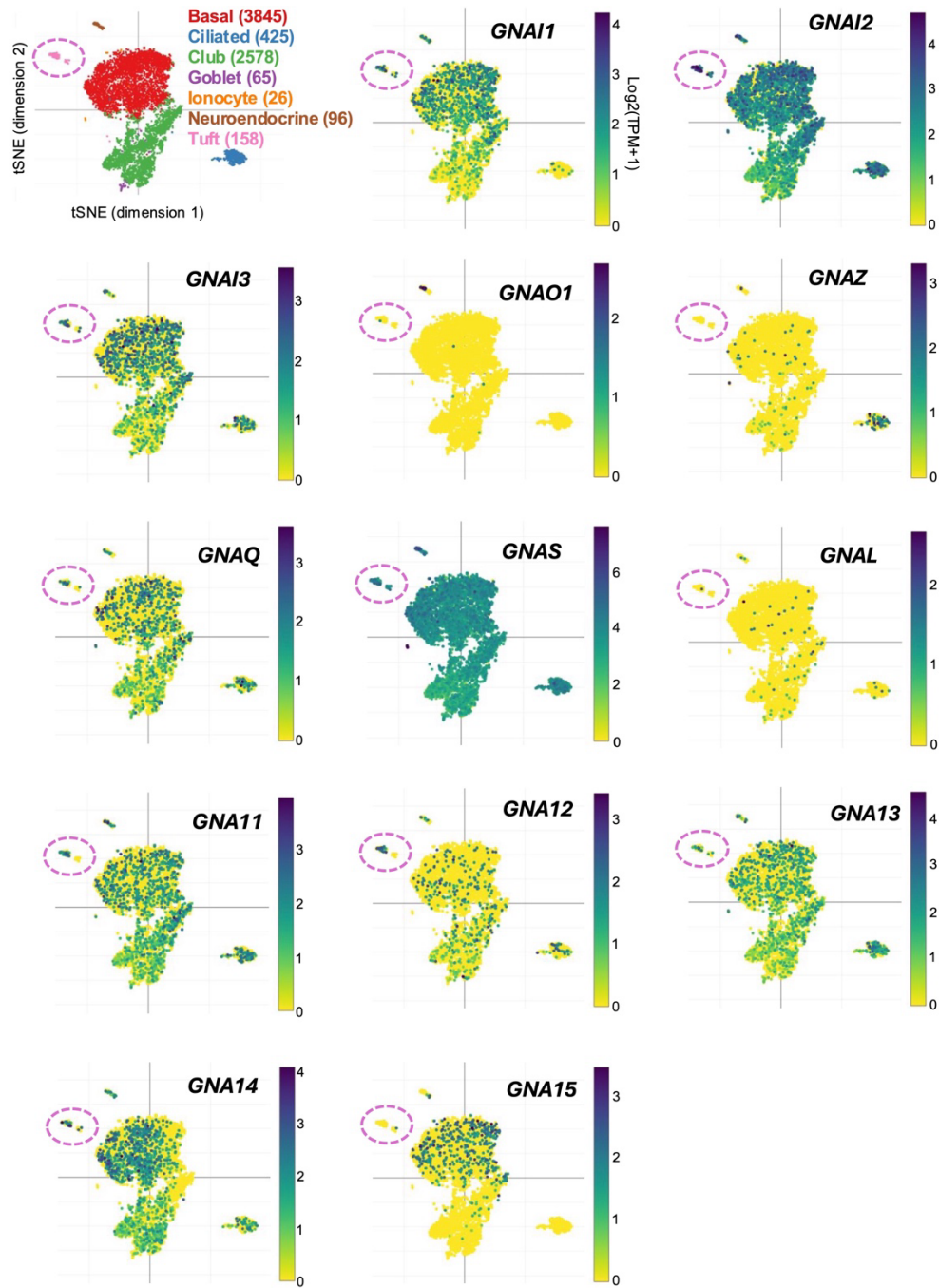

**Supplemental Figure 3.** tSNE plots of gene expression data from Montoro et al.,<sup>7</sup> showing expression of a variety of Gα subunits in mouse tracheal tuft cells, including members of the Gα<sub>i</sub> (*Gnai1*, *Gnai2*, and *Gnai3*), Gα<sub>q</sub> (*Gnaq* and *Gna11*), Gα<sub>s</sub> (*Gnas*), and Gα<sub>12/13</sub> (*Gna12* and *Gna13*) families. This suggests that tuft cells can activate multiple GPCR signaling pathways beyond gustducin. Data were accessed and plotted through the Broad Institute Single Cell Portal (https://singlecell.broadinstitute.org/single_cell/study/SCP163/airway-epithelium).

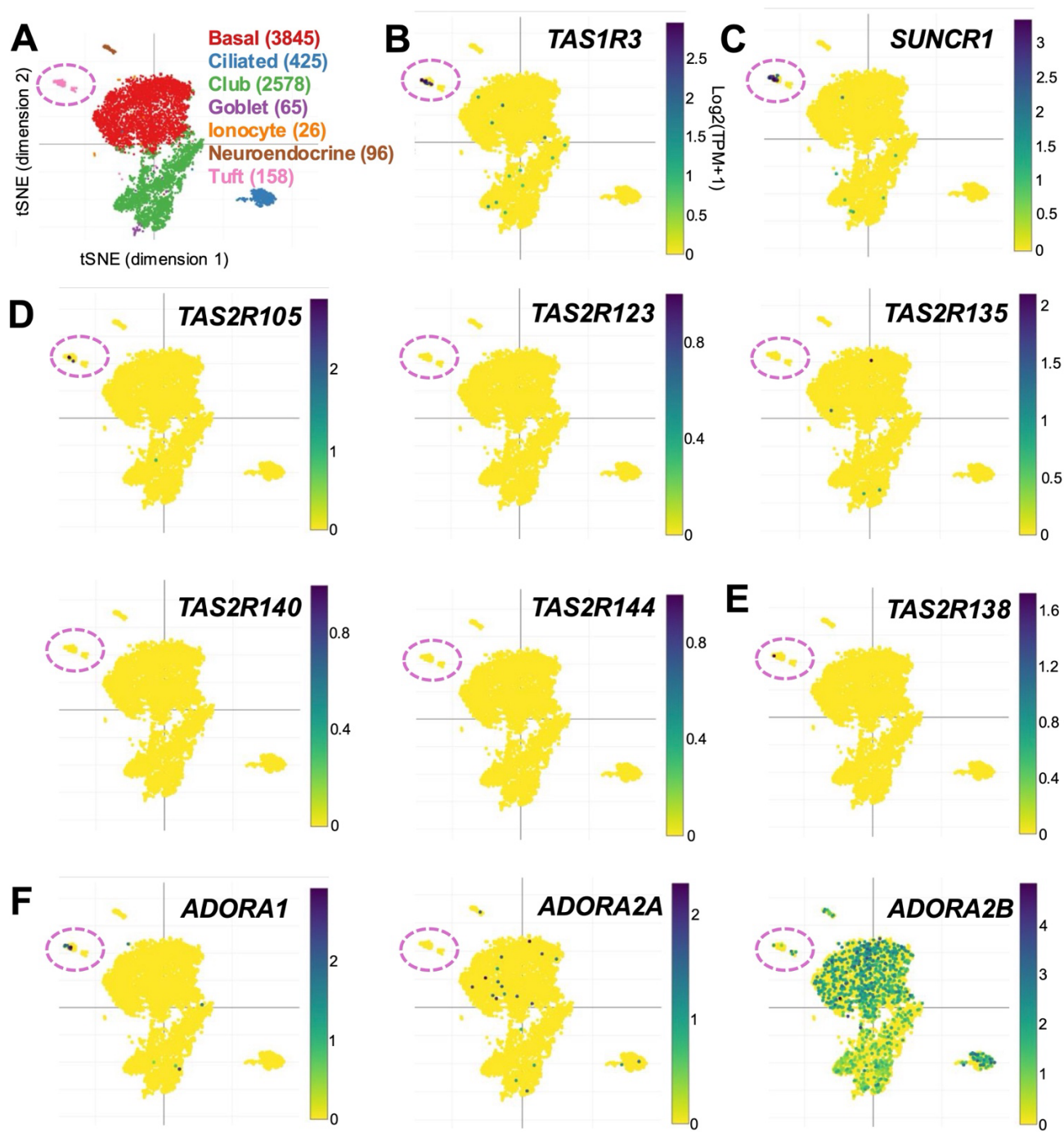

**Supplemental Figure 4.** tSNE plots of gene expression data from Montoro et al.,<sup>7</sup> showing expression and enrichment of GPCR genes in mouse tracheal tuft cells, including the sweet/umami taste receptor subunit *Tas1r3*, the succinate receptor *Sucnr1*, the bitter taste receptor *Tas2r105*, and the A1 adenosine receptor *Adora1*. Data were accessed and plotted through the Broad Institute Single Cell Portal (https://singlecell.broadinstitute.org/single_cell/study/SCP163/airway-epithelium).

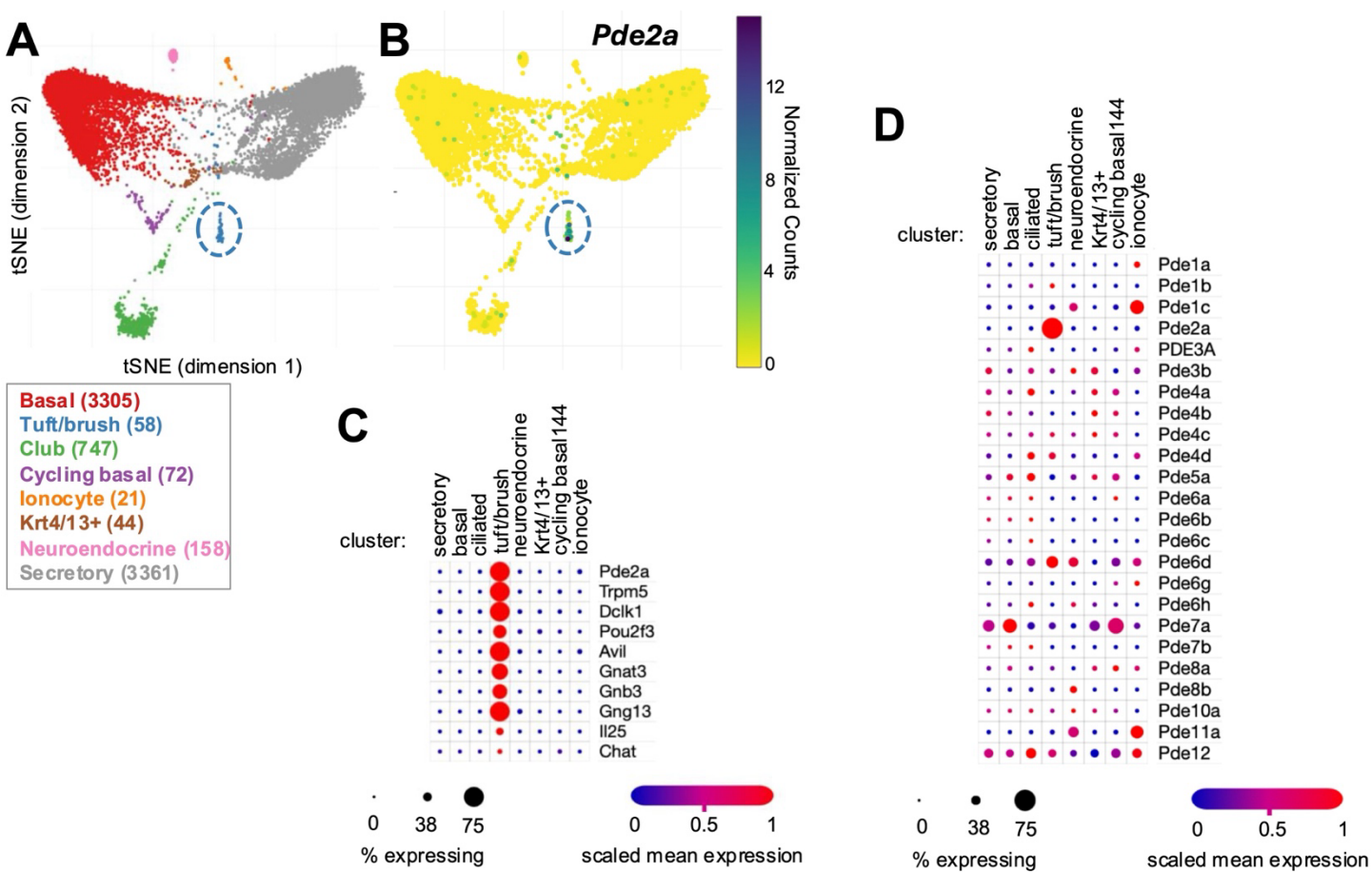

**Supplemental Figure 5.** Gene expression data from Plasschaert et al.,<sup>8</sup> supporting enrichment of *Pde2a* in mouse airway tuft cells. Data were accessed and plotted through the Broad Institute Single Cell Portal (https://singlecell.broadinstitute.org/single_cell/study/SCP64/a-single-cell-atlas-of-the-airway-epithelium-reveals-the-cftr-rich-pulmonary-ionocyte).

### Targeting cyclic-AMP/phosphodiesterase signaling to quiet nasal tuft cell calcium responses

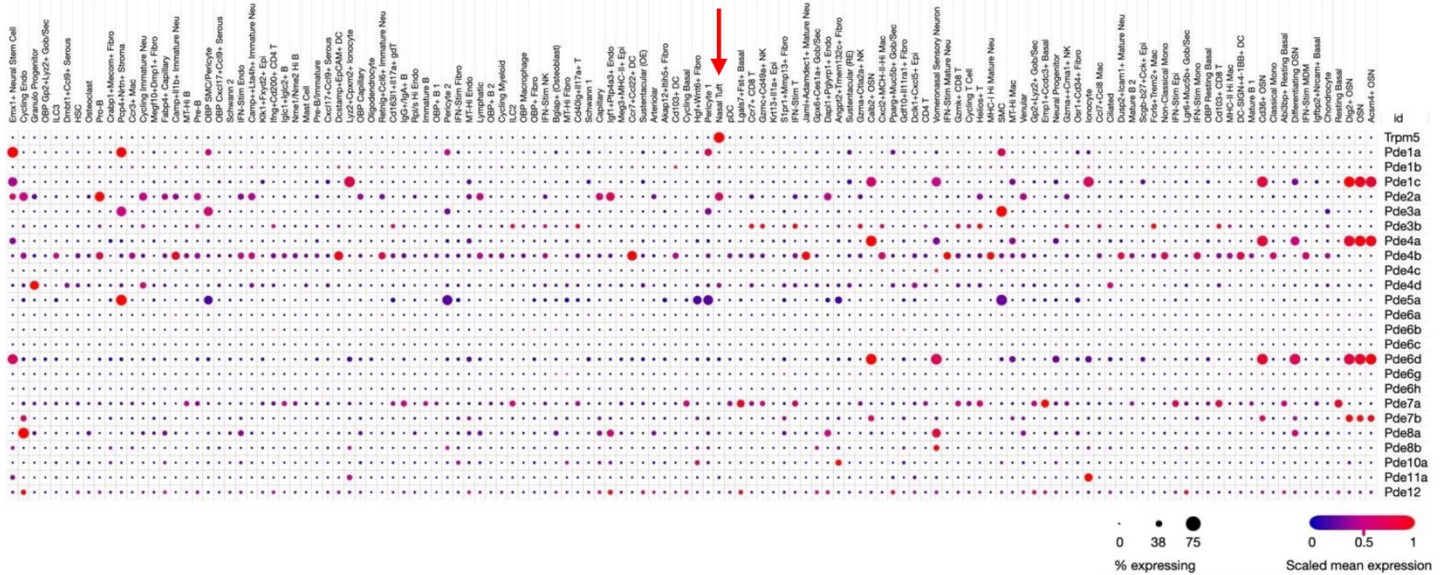

**Supplemental Figure 6.** Gene expression data from Kazer et al.<sup>9</sup> of mouse nasal tissue showing expression of *Trpm5* and *Pde2a* in mouse nasal tuft cells. Data were accessed and plotted through the Broad Institute Single Cell Portal (https://singlecell.broadinstitute.org/single_cell/study/SCP2216/primary-nasal-viral-infection-rewires-the-tissue-scale-memory-response-primary-infection)

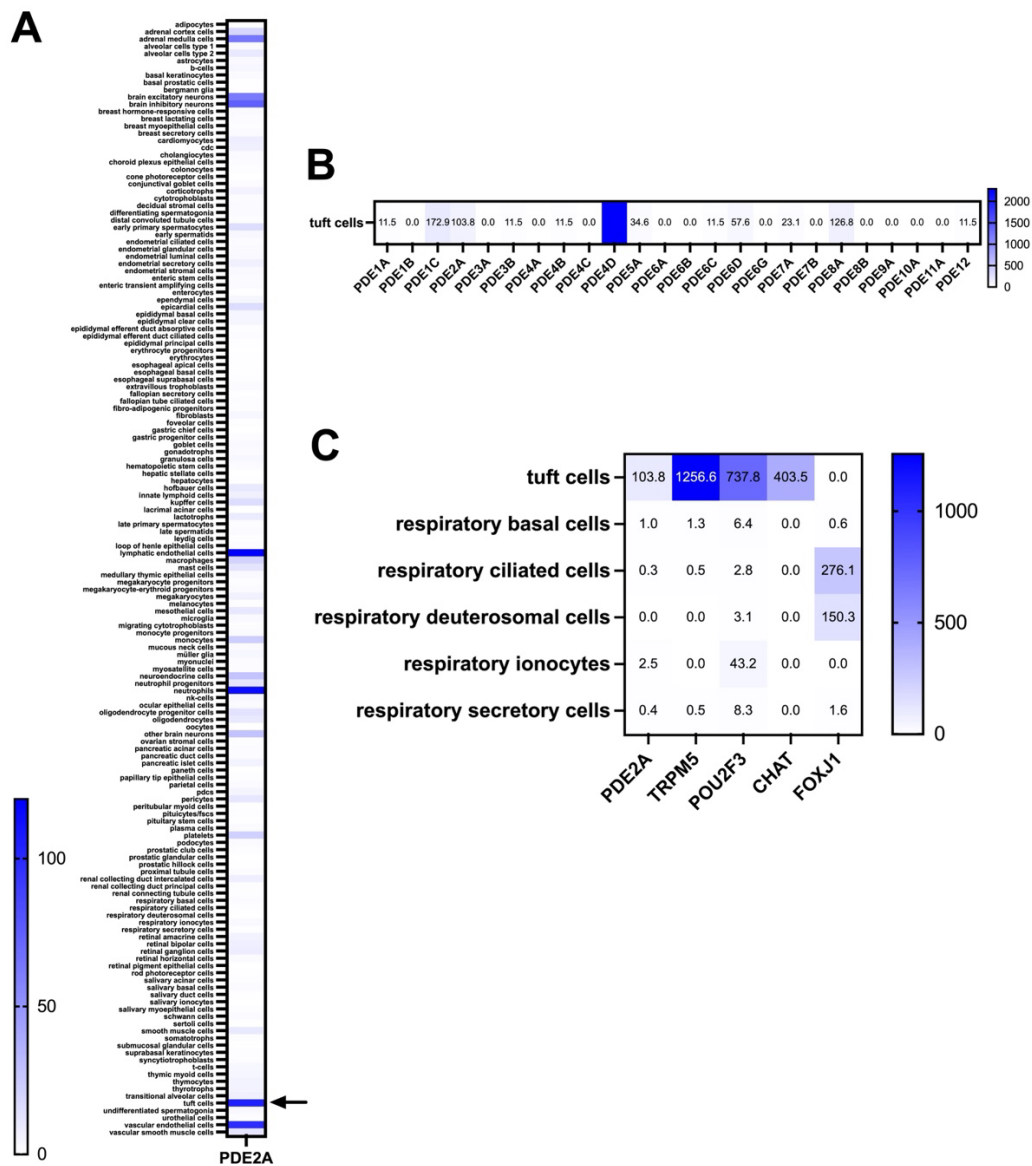

**Supplemental Figure 7.** PDE2A expression in human tuft cells. (A) Heat map showing expression of PDE2A in various human cell types from the Human Protein Atlas using single-cell RNAseq. This is consistent with a prior study that detected PDE2A expression in bulk human nasal mucosa.<sup>10</sup> (B) Heatmap showing PDE isoforms expression in human tuft cells. (C) Heatmap showing PDE2A expression in tuft cells compared with other respiratory epithelial cell types in the Human Protein Atlas. Tuft cell markers TRPM5, POU2F3, and CHAT, as well as the ciliated cell marker FOXJ1, are shown as controls. Graphs were made in GraphPad Prism from gene expression data downloaded from the Human Protein Atlas (https://www.proteinatlas.org/humanproteome/single+cell/single+cell+type/data#cell_type_data)

### Supplemental References

1. Lee RJ, Kofonow JM, Rosen PL, et al. Bitter and sweet taste receptors regulate human upper respiratory innate immunity. *J Clin Invest* 2014; 124:1393-1405.
2. Lee RJ, Chen B, Redding KM, et al. Mouse nasal epithelial innate immune responses to *Pseudomonas aeruginosa* quorum-sensing molecules require taste signaling components. *Innate Immun* 2014; 20:606-617.
3. Kuek LE, McMahon DB, Ma RZ, et al. Cilia Stimulatory and Antibacterial Activities of T2R Bitter Taste Receptor Agonist Diphenhydramine: Insights into Repurposing Bitter Drugs for Nasal Infections. *Pharmaceuticals (Basel)* 2022; 15.
4. Kouakou YI, Thompson JC, Tan LH, et al. Hops bitter beta-acids have antibacterial effects against sinonasal *Staphylococcus aureus* but also induce sinonasal cilia and mitochondrial dysfunction. *Int Forum Allergy Rhinol* 2024; 15:287-302.
5. Thompson JC, Park A, Kouakou YI, et al. Mechanisms of Mitochondrial Toxicity and Cytotoxicity Caused by *Pseudomonas aeruginosa* Pyocyanin in Human Nasal Epithelial Cells. *Int Forum Allergy Rhinol* 2026; 16:451-466.
6. Sisson JH, Stoner JA, Ammons BA, et al. All-digital image capture and whole-field analysis of ciliary beat frequency. *J Microsc* 2003; 211:103-111.
7. Montoro DT, Haber AL, Biton M, et al. A revised airway epithelial hierarchy includes CFTR-expressing ionocytes. *Nature* 2018; 560:319-324.
8. Plasschaert LW, Zilionis R, Choo-Wing R, et al. A single-cell atlas of the airway epithelium reveals the CFTR-rich pulmonary ionocyte. *Nature* 2018; 560:377-381.
9. Kazer SW, Match CM, Langan EM, et al. Primary nasal viral infection rewires the tissue-scale memory response. *bioRxiv* 2024; 18:2023.05.11.539887.
10. Saraiva LR, Riveros-McKay F, Mezzavilla M, et al. A transcriptomic atlas of mammalian olfactory mucosae reveals an evolutionary influence on food odor detection in humans. *Sci Adv* 2019; 5:eaax0396.
